# Leaf reflectance can surrogate foliar economics better than physiological traits across macrophyte species

**DOI:** 10.1101/2020.06.03.131375

**Authors:** Paolo Villa, Rossano Bolpagni, Monica Pinardi, Viktor R. Tóth

## Abstract

Macrophytes are key players in aquatic ecosystems diversity, but knowledge on variability of their functional traits, among and within species, is still limited. Remote sensing is a high-throughput, feasible option for characterizing plant traits at different scales, provided that reliable spectroscopy models are calibrated with congruous empirical data.

We sampled leaves from six floating and emergent macrophyte species common in temperate areas, covering different phenological stages, seasons, and environmental conditions, and measured leaf reflectance (400-2500 nm) and leaf traits (dealing with photophysiology, pigments and structure). We explored optimal spectral bands combinations and established non-parametric reflectance-based models for selected traits, eventually showing how airborne hyperspectral data can capture spatial-temporal macrophyte variability.

Our key finding is that structural - leaf dry matter content, leaf mass per area - and biochemical - chlorophyll-a content and chlorophylls to carotenoids ratio - traits can be surrogated by leaf reflectance with relative error under 20% across macrophyte species, while performance of reflectance-based models for photophysiological traits depends on species.

This finding shows the link between leaf reflectance and leaf economics (structure and biochemistry) for aquatic plants, thus supporting the use of remote sensing for enhancing the level of detail of macrophyte functional diversity analysis, to intra-site and intra-species scales.

## 1. Introduction

Ecotones laying at the interface between land and inland waters are among the most important (Tranvik *et al*., 2009; Mitsch *et al*., 2013), most productive (Bridgham *et al*., 2006) and most diverse (Keddy *et al*. 2009) ecosystems in temperate areas. The origin of such exceptional diversity is attributed to their transitional status, connecting aquatic and terrestrial biomes (Wetzel, 1990), coupled with a high spatial and temporal variability of environmental conditions (Spence, 1982).

Within this frame, aquatic plants, i.e. macrophytes, are key players, as they are hotspots of biogeochemical cycling, actively impact on ecosystem by regulating water flow and sedimentation, and promote biodiversity by attracting and sheltering large number of species (Schindler & Scheuerell, 2002; Franklin *et al*., 2008;Jeppesen *et al*., 2012; Wang *et al*., 2017). Adaptations to environmental heterogeneity is particularly evident for macrophytes, whose genetic variability and morpho-functional plasticity result in specific phenotypic and phenological patterns within the local population, shaping communities (Elzinga *et al*., 2007; Chuine, 2010; Tóth & Szabó, 2012; Tóth & Vári, 2013) and even individuals (de Kroon *et al*., 2005; Tóth & Palmer, 2016).

As multi-dimensional trait variability is a recent object of investigation in functional ecology (Messier *et al*., 2010; Hu *et al*., 2015; Butler *et al*., 2017), knowledge about ranges and interconnections of phenotypic and phenological variability within a community (population) is still relatively limited, all the more when aquatic plants are considered, due to their peculiar features and the environmental patchiness of their habitat, showing high abiotic and biotic variability (Vivian-Smith, 1997). The issue of disentangling the effects of inter-specific and intra-specific trait variability is at the centre of the current debate (Albert *et al*., 2012; Osnas *et al*., 2018), especially for key traits related to the leaf economics spectrum (LES; Wright *et al*., 2004). Moreover, spatial patterns of trait variability at local scale have been studied only by a limited number of works (Kumordzi *et al*., 2015; Sandel & Low, 2019) and some light has still to be shed on this level of heterogeneity in plant communities, with implications on productivity and connected processes (Funk *et al*., 2016).

Exploring this fine scale variability with direct measurements, usually carried out *in situ* and in laboratory, is very time and resource consuming, and often logistically constrained in aquatic systems. The level of maturity in platforms and techniques achieved by remote sensing (RS) make it a feasible and potentially very effective way forward in characterizing selected plant trait variability within communities at different geographic scales (Asner *et al*., 2015; Abelleira Martinez *et al*., 2016; Gamon *et al*., 2019; Dalla Vecchia *et al*., 2020), overcoming logistic and economic constraints.

In particular, the last two decades have seen the development of a range of applications for RS of plant biophysical and bio-chemical traits, with an intensification of this trend in last decade (Homolová *et al*., 2013; Verrelst *et al*., 2015; Wang & Gamon, 2019). RS-based works have by a large majority focused on terrestrial plants, from forest and grassland biomes (e.g Asner & Martin, 2008; Asner *et al*., 2015; Singh *et al*., 2015; Schweiger *et al*., 2017; Kattenborn *et al*., 2018; Serbin *et al*., 2019), but some recent developments on aquatic vegetation have been recently documented (Stratoulias *et al*., 2015; Villa *et al*., 2017). For being effective, applications of RS for mapping plant functional traits require an analysis of which traits can be modelled from spectral reflectance, and this is preferably done using data covering natural trait heterogeneity. Empirical approaches for assessing reflectance spectra as proxy of plant traits, at leaf or canopy level, normally employ spectroscopy-based methods ranging from parametric regression models input with spectral indices, computed as band combinations (e.g. Le Maire *et al*., 2008; Yao *et al*., 2010; Ali *et al*., 2017) to non-parametric regression models, such as partial least-square regression (e.g. Asner & Martin, 2008; Feilhauer *et al*., 2010; Ely *et al*., 2019; Fu *et al*., 2020).

Towards meaningful applications of RS for plant functional ecology, the fundamental question is: which leaf traits, within a specific plant group, can be reliably modelled (and which other cannot) using spectroscopy? With this study, we aim to provide an answer to this question for what concerns floating and emergent aquatic plants common in temperate areas, based on empirical data of leaf spectra and a set of leaf traits - dealing with photophysiology, pigments and leaf structure – collected *in situ* from six macrophyte species, under different times, seasons and environmental conditions, over three sites located in Europe.

The objectives of this work are: i) to assess the variability in leaf functional traits within and among macrophyte species, with attention to autochthonous *vs*. allochthonous taxa dualism; ii) to evaluate which functional traits – photophysiological, biochemical and structural – can be effectively modelled across macrophyte species using leaf reflectance, and which wavelength ranges and combinations are more sensitive to specific traits; and iii) to test how leaf reflectance-traits relations can be exploited using remote sensing data for providing information on spatial and temporal functional variability of macrophyte communities at inter- and intra-species level.

## 2. Materials and Methods

### 2.1. Sampling site descriptions

Macrophyte samples were collected from three temperate shallow lakes surrounded by wetlands and hosting abundant macrophyte communities, located in central and southern Europe: Lake Hídvégi (Hungary), Mantua lakes system (Italy), and Lake Varese (Italy).

Lake Hídvégi, located in western Hungary (46°38′ N, 17°08′ E; 110 m a.s.l.), is an artificial lake system built in 1985 as a part of the Kis-Balaton Water Protection System, that has the overall function of retaining inorganic nutrients and total suspended solids carried by the Zala River into Lake Balaton (Korponai *et al*., 2010). Lake Hídvégi is a shallow, eutrophic to hypertrophic, predominantly open-water habitat (area: 18 km^2^, average depth: 1.1 m) partly covered by floating macrophytes, with *Trapa natans* L. as dominant species and some presence of *Nuphar lutea* (L.) Sm. and *Nymphaea alba* L. (Dinka *et al*., 2008; Villa *et al*., 2017). Helophyte communities in the littoral zone are composed of reed beds (*Phragmites australis* (Cav.) Trin. ex Steud.) and cattail beds (*Typha* spp.).

Mantua lakes system, located in northern Italy plain (45°10′ N, 10°47′ E; 15 m a.s.l.), is composed by three dimictic shallow fluvial-lakes (area: 6.1 km^2^; average depth: 3.5 m), with two connected wetlands (upstream and downstream of lakes). The Superior, Middle and Inferior lakes are characterized by high turbidity and eutrophic conditions, and water level is kept fixed to prevent flooding the city of Mantua. Macrophyte communities in the system are populated by both autochthonous (*T. natans, N. lutea, N. alba* in open water areas, and *P. australis* in wetlands) and allochthonous species (Pinardi *et al*., 2011; Villa *et al*., 2017; Tóth *et al*., 2019): *Nelumbo nucifera* Gaertn., introduced into the lake around a century ago, and *Ludwigia hexapetala* (Hook. & Arn.) Zardini, H.Y. Gu & P.H. Raven, which started spreading here during the last decade.

Lake Varese, located in subalpine northern Italy (45°49′ N, 8°44′ E; 238 m a.s.l., is a monomictic, eutrophic lake (area: 14.2 km^2^; average depth: 10.9 m) subject to high anthropic pressures and nutrient loads. The southern shores of the lake host extensive stands of floating macrophytes, mainly *T. natans* and *N. lutea*, with some presence of *N. alba*. (Villa *et al*., 2015). Some tracts of littoral zone have been colonized in the last couple of decades by allochthonous species, *N. nucifera* and *L. hexapetala*.

### 2.2. Field measurements

Boat-based surveys were carried out in the three sites for three years (2016-2018), covering different times within the macrophyte growing season, spanning from late May to late July. Due to logistic and technical constraints, macrophyte beds sampled were not always the same ones in different years. During the surveys, leaf samples from 6 species showing sensitive presence in the study areas (*L. hexapetala, N. nucifera, N. alba, N. lutea. P. australis, T. natans*) were measured and collected over various locations, to incorporate intra-site variability. A summary of sampling locations for each species and survey date is provided in Supplementary Table 1. Leaf samples, either floating or emergent above water, were collected from plants growing in dense (canopy fraction cover > 60%) and homogeneous stands, within 3 m of the water edge; from each plant sampled, the youngest, mature leaf directly exposed to sunlight was taken for measurements.

#### 2.2.1. Leaf spectral reflectance

Leaf reflectance in the visible to shortwave infrared range (350 – 2500 nm, spectral resolution of 3 nm for wavelengths under 1000 nm, and < 8 nm up to 2500) was measured using a portable full range spectroradiometer (SR-3500, Spectral Evolution, Lawrence, USA), following the protocol described in Tóth *et al*. (2019). After the 20-minute dark adaptation, leaves were de-shadowed and laid on a flat neoprene plate (reflectance factor < 5%) in order to minimize background reflection of light transmitted through the leaves. Leaf reflected radiance was measured at contact using a probe equipped with 5-watt internal light source under near-steady state conditions, i.e. 60 seconds after de-shadowing. Each spectrum is the result of 10 averaged scans, and automatic integration time optimization was used, with a maximum allowed of 50 ms per scan. Leaf spectra were finally calibrated to reflectance using reflected radiance from a Spectralon panel (Labsphere, North Sutton, USA; reflectance factor > 99% for wavelengths under 1500 nm, and > 95% up to 2500 nm) as reference.

#### 2.2.2. Leaf photophysiology

Photophysiological traits of macrophytes were assessed using chlorophyll fluorescence measured with a PAM-2500 chlorophyll fluorometer (Heinz Walz GmbH, Germany) over the same leaves sampled for spectral reflectance measurements. Data were collected occasionally between 09:00 and 15:00, standard solar time. Relevant fluorescence yield data (initial fluorescence yield - F_0_, maximal fluorescence yield - F_m_) were measured following the protocol described in Tóth *et al*. (2019) on mature, healthy-looking leaves after a dark-adapting period of 20 minutes with a pulse of a saturated light (630 nm, intensity 3000 μmol m^−2^ s^−^1). Photochemical PSII efficiency (F_v_/F_m_), coefficient of photochemical quenching (qP), coefficient of non-photochemical quenching (qN), maximum electron transport rate of the photosystem II (PSII) (ETR_max_), theoretical saturation light intensity (I_k_), and maximum quantum yield for whole chain electron transport (α) were calculated using fluorescence yield data and a light response curve - function of ETR to photosynthetically active radiation (PAR) in 11 steps between 5 and 787 μmol m^−2^ s^−1^ - described by an exponentially saturating equation (Eilers & Peeters, 1988; Genty *et al*., 1989). Leaf absorbance was set at 0.84 for calculation of ETR for all the species sampled.

#### 2.2.3. Leaf biochemical and structural traits

Leaf pigments, as biochemical traits directly connected with photosynthetic apparatus, and leaf structural traits were measured on our samples to represent foliar economics expressing the inner trade-off between resource availability and structural investments, which lies at the core of the LES.

Two leaf discs (0.6 cm in diameter) were cut with a cork borer from each fresh leaf, in the vicinity of where chlorophyll fluorescence was measured. Disks were stored in aluminium foil at sub 0 temperature in a camping fridge until they were transferred to the laboratory, i.e. within maximum 4 hours.

Half of the discs sampled were stored in −20° C freezer, to be used for pigments extraction, following the protocol described in Tóth *et al*. (2019). Upon extraction, they were homogenised in liquid N_2_ in a grinder, subsequently extracted in acetone solution (80%), and stored in a fridge overnight. The extracts were centrifuged and the supernatant collected and stored at –20 °C. The full spectra of absorbance of the extracts were measured between 400 nm and 750 nm using a spectrophotometer (Shimadzu UV-2401PC, dual-beam), at 1 nm resolution. Finally, pigment concentrations, i.e. chlorophyll-a (Chl-a), chlorophyll-b (Chl-b), and total carotenoids (Car), were calculated using empirical formulae (Wellburn, 1994) and reported on leaf area basis (µg cm^-2^). Pigments ratios were then calculated, as Chl-a to Chl-b ratio (Ca/Cb) and total chlorophylls (Chl-a + Chl-b) to total carotenoids ratio (Chl/Car).

The other half of leaf discs were weighted with a precision balance (Mettler Toledo AB104, 0.0001 g accuracy) to measure their fresh weight, and were then dried in ventilated oven at 70° C for 48h, after which the dry weight was measured using the same balance. From these data, dry matter content (DMC) and leaf mass per area (LMA) of each disc were calculated as the ratio of dry weight to fresh weight (g g^-1^), and the ratio of dry weight to disc area (g m^-2^) respectively.

#### 2.2.4. Interspecific and intraspecific variability

Variability in traits across species was tested using non parametric Kruskal-Wallis One Way ANOVA, and pairwise comparisons were performed using *post-hoc* Dunn’s test, with p-value adjustment computed according to Benjamini-Hochberg method. Intraspecific variability of selected traits was assessed by calculating the coefficient of variation (CV) of each trait for every species, using only leaf samples collected at peak of growth (i.e. July) to reduce differences due to phenology.

### 2.3. Spectral indices and leaf traits

For each possible combination of two spectral reflectance bands measured as described in Section 2.2.2 within the range 400-2500 nm (987 bands), the normalized difference spectral index (NDSI) was calculated using a custom-made R script (Stratoulias *et al*., 2015). NDSI are frequently used in RS because they offer the advantage of summarizing spectra information, while reducing uncertainty due to sensor differences and atmospheric effects and bias due to differences in vegetation background (Blackburn, 2006; Glenn *et al*., 2008; Ustin *et al*., 2009), and are defined as:

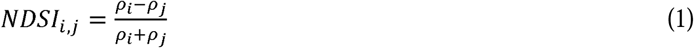

Where _ρ_ is leaf reflectance measured at spectral bands *i* or *j*.

Using the same R script, the correlation (Pearson’s *r*) between each leaf parameter, i.e. photophysiology, pigments and structural trait, and every calculated NDSIs was calculated together with the corresponding p-value (p). Correlation plots featuring the coefficient of determination (R_cal_^2^), calculated as the square of *r*, between each leaf parameter and the complete combinations of NDSIs derived from leaf reflectance were drawn for significant relations (p < 0.01), in order to highlight optimal two-band combinations proxies of investigated leaf photophysiology (α, ETR_max_, I_k_, F_v_/F_m_, qN, qP), pigments pool (Chl-a, Chl-b, Car, Ca/Cb, Chl/Car), and structural traits (DMC, LMA).

### 2.4. Hyperspectral modelling of leaf traits

Partial least-square regression (PLSR) modelling (Geladi & Kowalski, 1986) was used to further explore the relations between leaf reflectance and selected leaf traits across floating and emergent macrophyte species, using package *pls 2.7*, implemented in R (Mevik & Wehrens, 2007; R Development Core Team, 2018). PLSR is a powerful tool for modelling vegetation parameters using spectral datasets, because it is suitable for cases when number of predictors is larger than that of observations and can handle multi-collinearity in predictor variables, as frequently happens with narrow-band hyperspectral data (Wold *et al*., 2001; Feilhauer *et al*., 2010).

In order to focus on meaningful relations, PLSR models were calibrated only for leaf traits that showed a minimum sensitivity to spectral reflectance, that is scoring a R_cal_^2^ against all combinations of NDSIs higher than 0.15 (with p < 0.01), while simultaneously excluding mutually correlated traits (i.e. scoring Pearson’s *r vs*. any other trait < −0.5, or > 0.5). The number of PLSR components used for each calibrated model (with 30 components set as maximum) was optimized through minimization of the root mean square error of prediction (RMSEP) via leave-one-out cross-validation (LooCV). The best model for each trait was eventually selected by setting the number of PLSR components corresponding to minimum cross-validation RMSEP, and model performance was assessed comparing measured with PLSR predicted trait values through the coefficient of determination of measured vs. predicted traits via LooCV (R_CV_^2^). The relative importance of different wavelengths and spectral ranges to PLSR models calibrated for each variable was assessed computing the Variable Importance of Projection (VIP; Singh *et al*., 2015).

### 2.5. Airborne hyperspectral images

Airborne hyperspectral images of Lake Hídvégi and Mantua lakes system were acquired from the APEX imaging spectrometer (Schaepman *et al*., 2015) on 19 July 2014 (around 12:00 local time) and 27 September 2014 (around 13:45 local time), respectively. Visible to near-infrared range APEX data were processed, resulting in hyperspectral images composed by 98 spectral bands (425-905 nm), with 3-10 nm spectral resolution.

APEX data were calibrated to at-sensor radiance units and georeferenced based on sensor’s GPS/IMU at 5 m spatial resolution on the ground. Surface reflectance was finally derived by applying atmospheric correction based on MODTRAN-4 code and optimized for water targets (De Haan *et al*., 1991).

To demonstrate the potential of hyperspectral remote sensing data in highlighting spatial patterns of macrophyte traits, maps of spectral proxies were derived from APEX surface reflectance bands by calculating the NDSI based on the optimal two-band combination for leaf traits whose NDSI-trait relation scored an R _cal_^2^ > 0.4 within the spectral range covered by APEX bands.

## 3. Results

### 3.1. Variability of macrophyte leaf traits

Regarding traits related to photosynthetic performance, allochthonous species showed significant differences with autochthonous ones, i.e. *Nelumbo* scored high α (adjusted p < 0.01 from all pairwise *post-hoc* Dunn’s tests; Supplementary Fig. S1), and *Ludwigia* scored high ETR_max_, I_k_ and qP (adjusted p < 0.001 from all pairwise *post-hoc* Dunn’s tests; Supplementary Fig. S1). Moreover, regulated thermal dissipation of excess absorbed light (qN) was found being slightly lower in *Nelumbo* and *Ludwigia*, as well as in *Trapa*, compared to nymphaeids (*Nuphar* and *Nymphaea*) and *Phragmites* (Fig. 1; Supplementary Fig. S1). *Phragmites* showed quantum efficiency of photosystem II (F_v_/F_m_) slightly higher than all other species, except *Nymphaea* (p < 0.05 from pairwise *post-hoc* Dunn’s tests; Supplementary Fig. S1).

**Figure 1.**
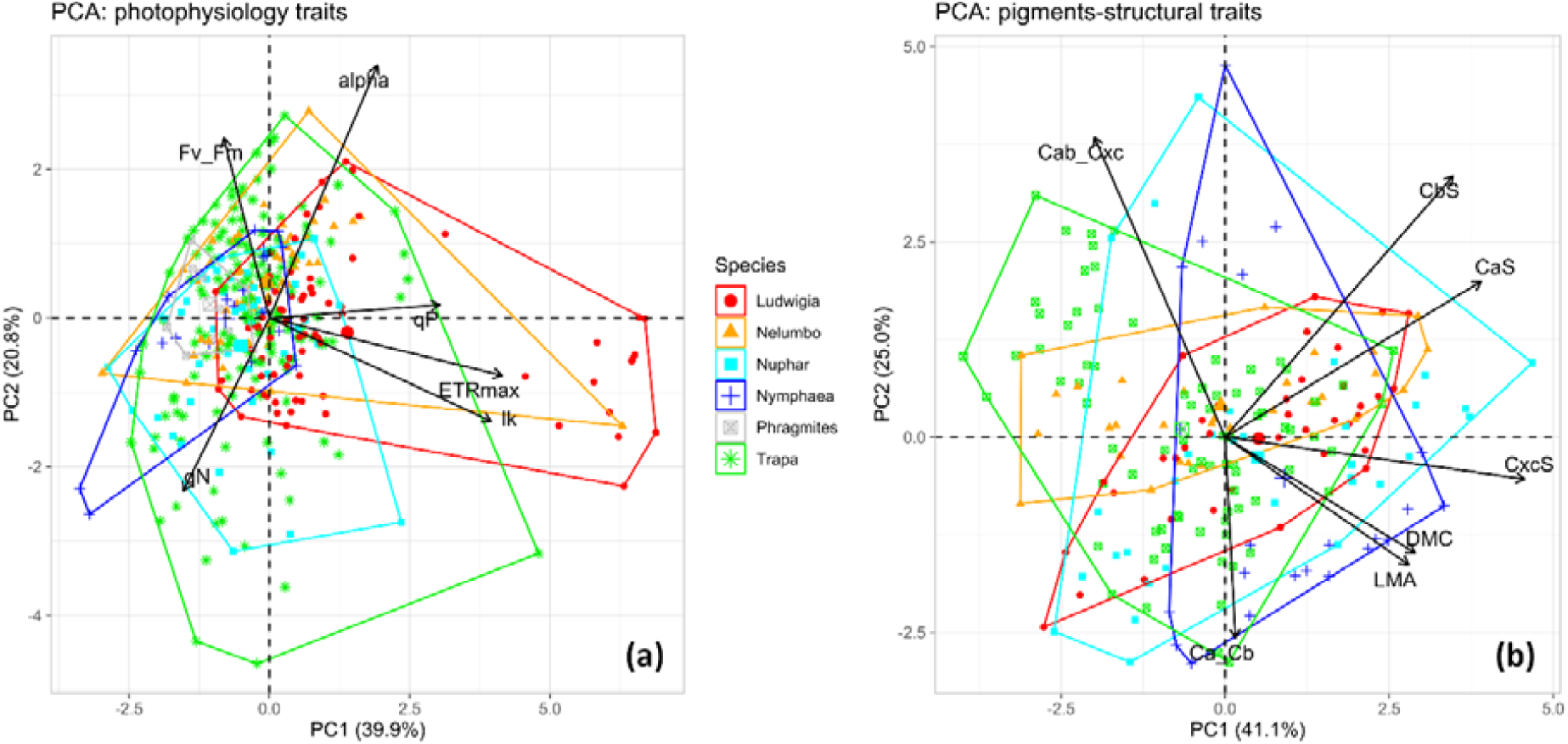
Bi-plots of the first two Principal Components of leaf photophysiological parameters (a) and foliar economics (b) expressed as pigments content and leaf structural traits, measured over all macrophyte species sampled. alpha: maximum quantum yield for whole chain electron transport; ETRmax: maximum electron transport rate for PSII; Ik: theoretical saturation light intensity for PSII; Fv_Fm: PSII photochemical efficiency; qP: coefficient of photochemical quenching; qN: coefficient of non-photochemical quenching; CaS: chlorophyll a content; CbS: chlorophyll b content; Cxc: total carotenoids content; Ca_Cb: chlorophyll a to chlorophyll b ratio; Cab_Cxc: chlorophylls to carotenoids ratio; DMC: leaf dry matter content; LMA: leaf mass per area.

*Ludwigia* displayed Chl-a values slightly higher than autochthonous species (p < 0.05 from all pairwise *post-hoc* Dunn’s tests; Supplementary Fig. S1), while no significant difference across species was observed for Chl-b, and lowest carotenoid content was found in *Nelumbo* and *Trapa* (p < 0.05 from all pairwise *post-hoc* Dunn’s tests; Supplementary Fig. S1). Nymphaeids got the lowest Ca/Cb ratio among species (p < 0.05 from pairwise *post-hoc* Dunn’s tests), with *Ludwigia* and *Trapa* occupying the high band of scores (Supplementary Fig. S1). A strong level of segmentation at species level was shown for Chl/Car ratio (p < 0.001, Kruskal-Wallis One Way ANOVA): *Nelumbo* and *Trapa* exhibited the highest scores (p < 0.001 from pairwise *post-hoc* Dunn’s tests) and *Ludwigia, Nuphar, Nymphaea* followed in decreasing Chl/Car (Supplementary Fig. S1).

Compared to autochthonous species, leaf DMC was higher (p < 0.001 from all pairwise *post-hoc* Dunn’s tests) and LMA lower (p < 0.001 from all pairwise *post-hoc* Dunn’s tests) for both *Ludwigia* and *Nelumbo* (Fig. 1; Supplementary Fig. S1). Inter-species DMC patterns tend to be similar to those of Chl/Car, with the exception of *Trapa*, and a strong differentiation in LMA was observed among species, with nymphaeids scoring the highest values and *Trapa* spanning the widest range (Fig. 1; Supplementary Fig. S1).

Leaf data collected at peak of growth (i.e. samples collected in July) exhibited a notable degree of intraspecific variability for most of the traits (Fig. 1), as maximum scores of CV across all species was larger than 0.29, with the only exceptions of _α_ and F_v_/F_m_ (Table 1). *Phragmites* presented the lowest variable set of photophysiology traits except for qP (CV < 0.19), while nymphaeids had the most variable traits (_α_, qP, Ca/Cb and Chl/Car for *Nymphaea*; Chl-a, Chl-b and Car for *Nuphar*). *Ludwigia* showed very high plasticity in photophysiology parameters ETR_max_, I_k_, and qN (CV > 0.32), and *Trapa* scored highest intraspecific ranges of DMC and LMA (CV > 0.29).

**Table 1.** Coefficient of variation (CV) of all leaf traits measured over macrophytes species sampled (highest scores across species for each trait are in bold, lowest in italic) at peak of growth conditions (July).

| Species | Photophysiology parameters |  |  |  |  |  | Leaf pigments |  |  |  |  | Leaf structure |  |
| --- | --- | --- | --- | --- | --- | --- | --- | --- | --- | --- | --- | --- | --- |
| | $\alpha$ | ETR<sub>max</sub> | I<sub>k</sub> | Fv/F<sub>m</sub> | qN | qP | Chl-a | Chl-b | Car | Ca/Cb | Chl/Car | DMC | LMA |
| *Ludwigia* | 0.134 | **0.886** | **0.695** | 0.060 | **0.325** | 0.111 | 0.303 | 0.295 | 0.243 | 0.098 | 0.123 | 0.160 | 0.163 |
| *Nelumbo* | 0.139 | 0.277 | 0.205 | 0.046 | 0.263 | 0.194 | 0.286 | 0.266 | 0.299 | 0.089 | 0.093 | 0.203 | 0.216 |
| *Nuphar* | 0.149 | 0.279 | 0.255 | 0.064 | 0.133 | 0.242 | **0.313** | **0.434** | **0.307** | 0.186 | 0.302 | 0.086 | 0.171 |
| *Nymphaea* | **0.202** | 0.396 | 0.244 | 0.029 | 0.183 | **0.378** | 0.190 | 0.368 | 0.293 | **0.293** | **0.363** | 0.136 | 0.217 |
| *Phragmites* | 0.074 | 0.172 | 0.187 | 0.044 | 0.072 | 0.156 |  |  |  |  |  |  |  |
| *Trapa* | 0.150 | 0.318 | 0.389 | **0.099** | 0.302 | 0.214 | 0.196 | 0.185 | 0.258 | 0.098 | 0.162 | **0.292** | **0.433** |

### 3.2. Spectral indices as proxies for macrophyte leaf traits

The correlations of every two-band combination NDSI correlations with investigated leaf traits measured on all macrophyte species sampled are highlighted in Fig. 2-4, showing the optimal spectral proxies for each trait in the visible to near-infrared (VNIR) spectral range (400-1000 nm) and in full spectral range (400-2500 nm), i.e. extending to shortwave infrared (SWIR) wavelengths.

**Figure 2.**
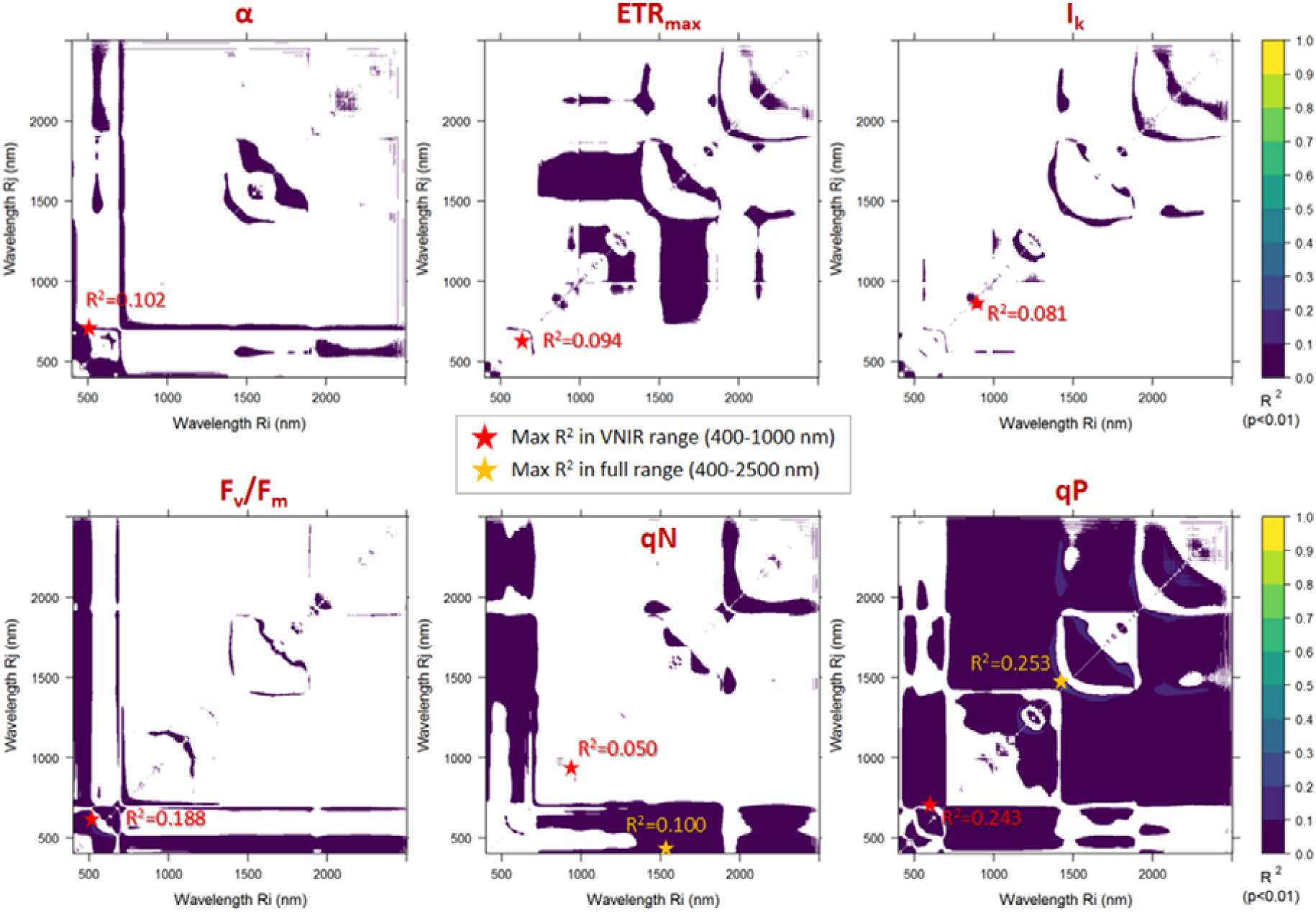
Statistically significant (p < 0.01) NDSI correlations with photophysiological parameters measured on all macrophyte species sampled (N = 324).

The heterogeneity of measured samples not only in terms of species, but also in terms of location (three study sites) and time of sampling (three years, different months) - which implies that our dataset includes a high variability in genetic features, seasonal cycles, stage of growths, plant conditions and environmental settings-overall resulted in relatively low correlation patterns (R_cal_^2^ < 0.26) between leaf reflectance derived NDSI and measured photophysiological traits (Fig. 2). In particular, negligible correlations with any NDSI combinations were shown across all species for _α_, ETR_max_, I_k_, qN (overall maximum R_cal_^2^ = 0.10, even if p < 0.01). Slightly higher but still moderately weak correlations are shown for F_v_/F_m_ and qP, with R _cal_^2^ peaking at 0.19 and 0.25, respectively (Fig. 2). The optimal NDSI for F_v_/F_m_ combined reflectance in the range of green visible light (524 nm, 581 nm), while for qP the best spectral combinations lay in the SWIR range, roughly around 1400-1500 nm.

Notwithstanding the abovementioned heterogeneity that biases the reflectance-photophysiology relations, correlations between pigments and NDSI computed from leaf reflectance across species are moderate to high, as peak R_cal_^2^ found was always higher than 0.3 (Fig. 3). In particular, good correlation between NDSIs and Chl-a were found, particularly in the red-edge range (maximum R_cal_^2^ = 0.43 for NDSI_775,740_). As Chl-b is highly correlated with Chl-a in our dataset (*r* = +0.80), optimal band combinations are found around the same range highlighted for Chl-a (795 nm, 740 nm), yet with slightly weaker correlation (maximum R _cal_^2^ = 0.37). Among pigments, Car showed the strongest correlations, peaking for NDSIs featuring band combinations in the SWIR range (maximum R_cal_^2^ = 0.57 for NDSI_1644,1720_), while correlations decreased if spectral range was limited to VNIR (maximum R_cal_^2^ = 0.37). Ca/Cb ratio showed the weakest correlations of all pigment traits, with a peak R_cal_^2^ = 0.31 in the SWIR range (for NDSI_2181,2241_). Chl/Car ratio is instead very well surrogated by NDSIs combining visible to SWIR reflectance (maximum R_cal_^2^ = 0.60 for NDSI_611,1892_); slightly lower correlations, yet still the highest among all pigments traits measured (maximum R_cal_^2^ = 0.55), were scored restricting the spectral range to VNIR range, where optimal NDSI for Chl/Car combined reflectance in 433 nm and 665 nm, which indeed roughly correspond to absorption peaks of chlorophyll-a (Ustin *et al*., 2009).

**Figure 3.**
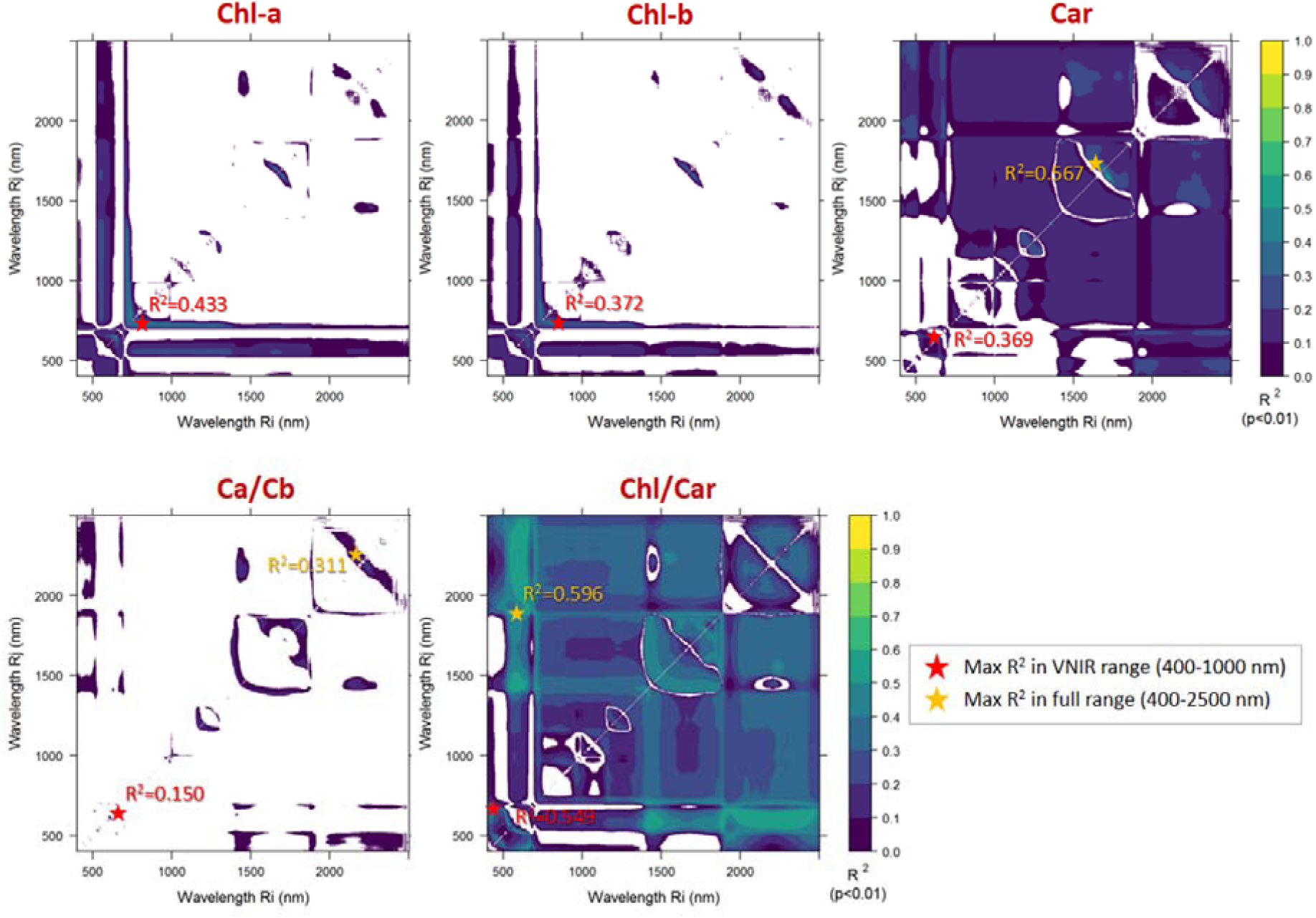
Statistically significant (p < 0.01) NDSI correlations with pigments content (on leaf area basis) and their balance measured on all macrophyte species sampled (N = 150).

Strong correlations were scored between NDSIs derived and leaf structural traits measured, connected to leaf economics, DMC and LMA (Fig. 4). DMC demonstrated a good sensitivity to leaf reflectance in the NIR to SWIR ranges, with R _cal_^2^ up to 0.47 (for NDSI_1196,1308_), and slightly weaker scores in the shorter VNIR wavelengths (maximum R_cal_^2^ = 0.35 for NDSI_929,941_). Among all leaf traits, LMA scored the highest correlations with NDISs, with a large subset of band combinations showing R_cal_^2^ > 0.6 in the NIR to SWIR ranges, and a peak R_cal_^2^ = 0.77 when two bands centred at 1415 and 2305 nm were used (Fig. 4).

**Figure 4.**
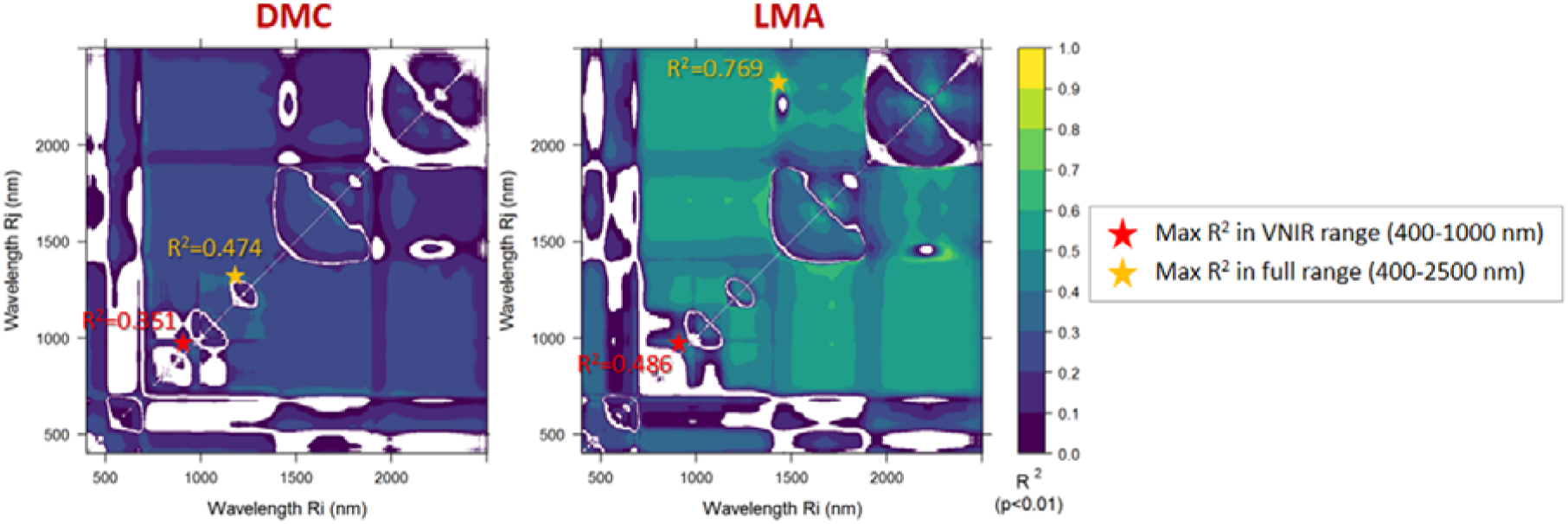
Statistically significant (p < 0.01) NDSI correlations with leaf economics traits, measured on all macrophyte species sampled (N = 152): leaf dry matter content (DMC) and leaf mass per area (LMA).

### 3.3. Leaf reflectance-traits relations

After having assessed mutual correlations of measured leaf traits, excluding the ones with |*r*| > 0.5 (Supplementary Fig. S8), PLSR models were calibrated for traits scoring non negligible correlations with best performing NDSIs presented in previous section (R_cal_^2^ > 0.15): F_v_ /F_m_, qP, Chl-a, Ca/Cb, Chl/Car, DMC, LMA.

The best fit PLSR models for F_v_/F_m_ and qP require 11 components, with RMSEP of 0.053 and 0.125 respectively (Supplementary Fig. S9). Matching between measured and predicted photophysiology traits was low to moderate for F_v_/F_m_ (R_CV_ ^2^ = 0.21) and qP (R_CV_ ^2^ = 0.33), even if estimation error seem to be reducing when extreme values - probably due to stress conditions non visibly detected when leaves were chosen for sampling – are excluded, i.e. for F_v_/F_m_ > 0.65 and for 0.4 < qP < 0.8 (Fig. 5).

**Figure 5.**
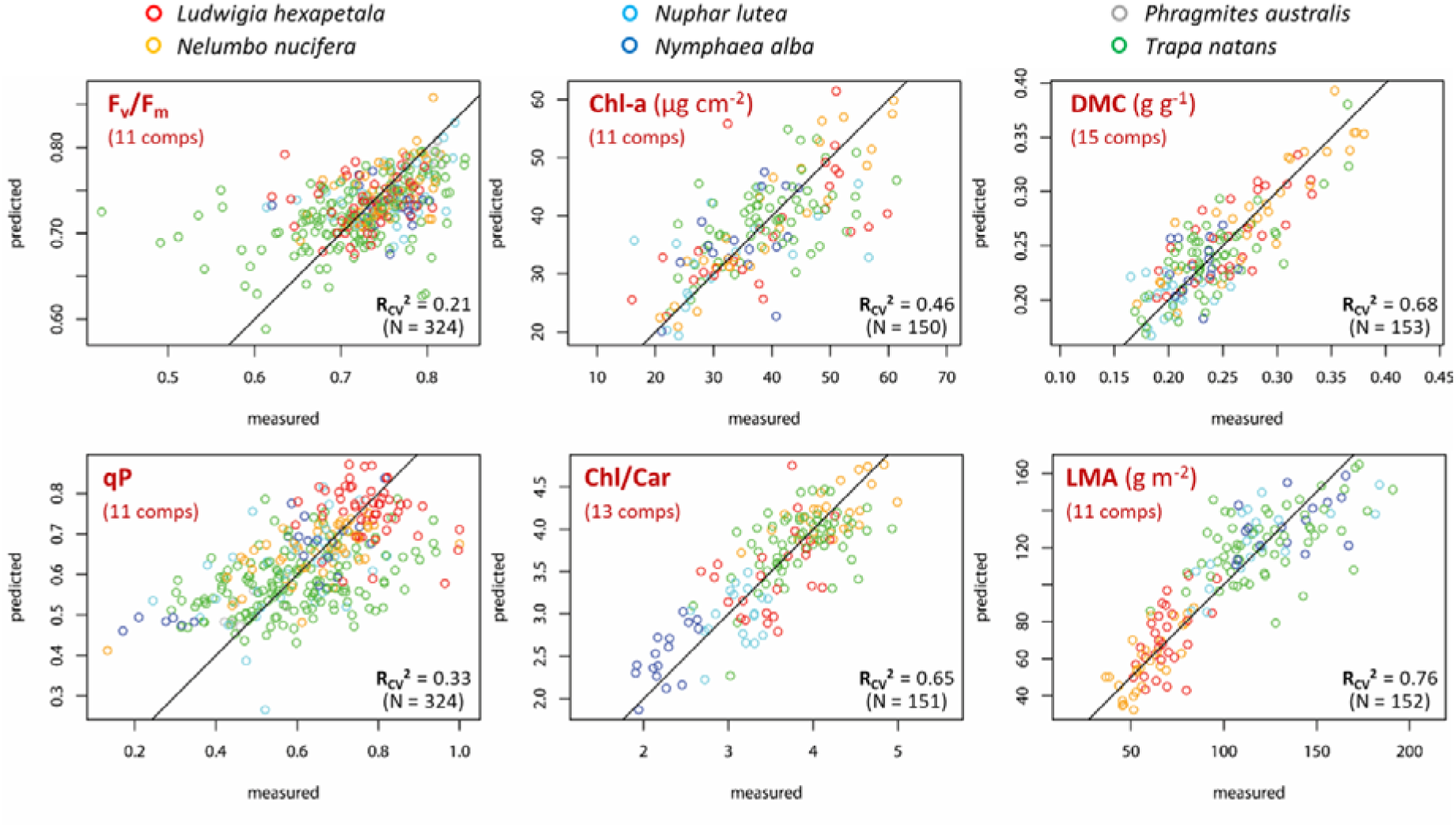
Comparison of leaf traits measured and predicted by best fit PLSR models for key leaf traits of selected macrophyte species, computed via leave-one-out cross-validation.

PLSR models based on leaf reflectance are quite effective in predicting macrophyte Chl-a content (on area basis) and Chl/Car ratio across species, with best fit models requiring 11 and 13 components (Fig. 5) respectively, achieving RMSEP of 7.52 µg cm^-2^ (R _CV_^2^ = 0.46) and 0.393 (R _CV_^2^ = 0.65). Fig. 6 shows that VIP scores are high around 550-560 nm and 705-710 nm for Chl-a model and around 705-710 nm and 1400 nm for Chl/Car model. Conversely, modelling performance for Ca/Cb is the lowest among traits considered here (R _CV_ ^2^ = 0.20), suggesting that spectral reflectance might not be a good proxy for this trait, at least for macrophyte species.

**Figure 6.**
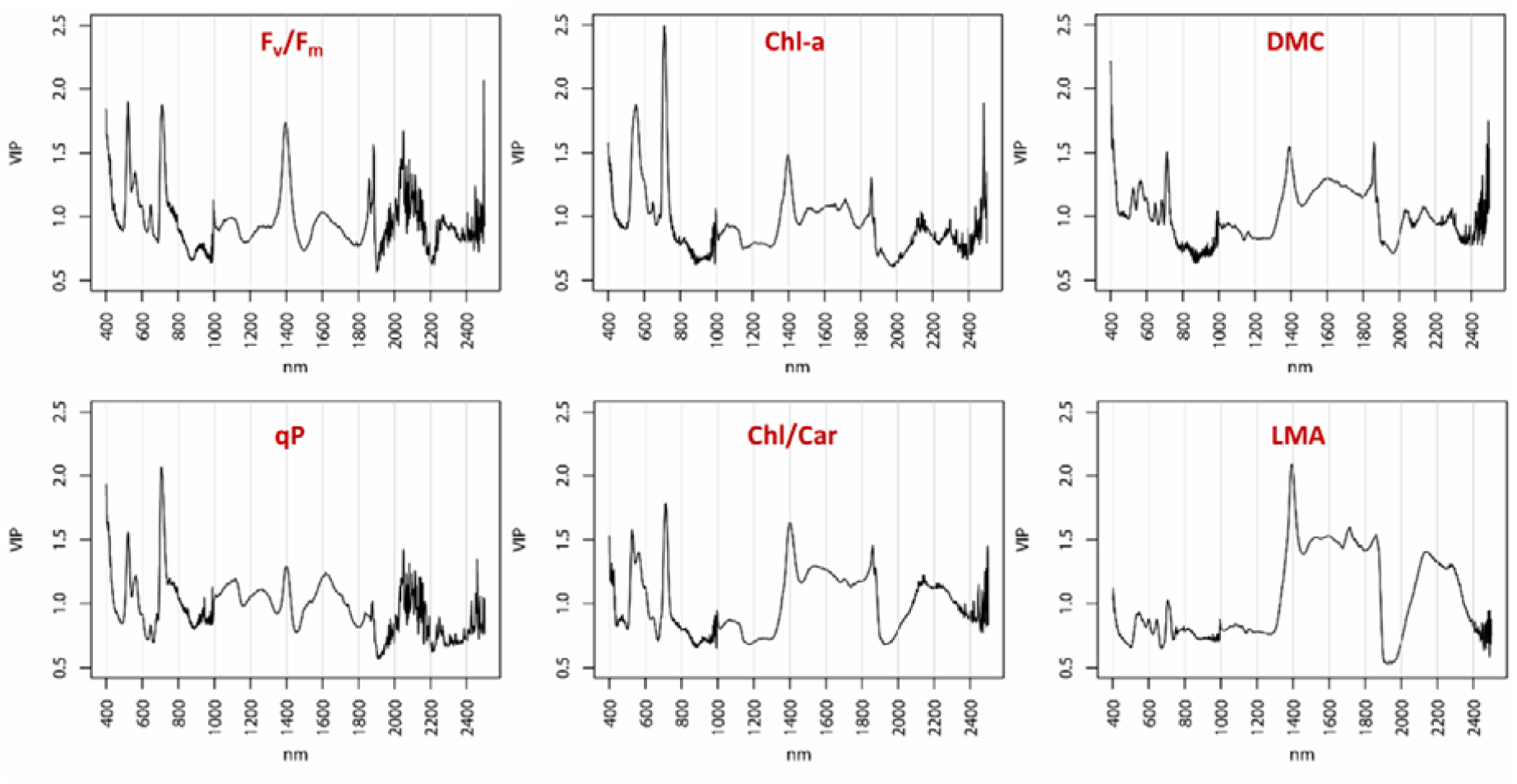
Variable Importance of Projection (VIP) scores of PLSR models of key leaf traits of selected macrophyte species, computed for each wavelength over.

DMC and LMA are the leaf traits better surrogated by reflectance-based models (Fig. 5, Supplementary Fig. S9), with matching between measured and PLSR predicted scores up to RMSEP = 0.026 g g^-1^ (i.e. 2.6% leaf weight) for DMC (R _CV_^2^ = 0.68 with 15 components) and up to RMSEP = 18.6 g m^-2^ for LMA (R _CV_^2^ = 0.76 with 11 components), across the whole range of measures in our dataset. VIP scores for DMC model are high around 400-410 nm and 1390-1400 nm, while for LMA model the wavelengths in VNIR range do not contribute much to prediction and peak VIP are scores between 1400 and 1850 nm, as well as between 2120 and 2300 nm (Fig. 6).

### 3.4. Mapping macrophyte traits from airborne hyperspectral images

The potential for application of the main findings described in previous sections was demonstrated by producing spatial variability maps of selected macrophyte traits through specific NDSIs highlighted in section 3.2 calculated from airborne hyperspectral data. Airborne data were collected over Lake Hídvégi and Mantua lakes system from APEX during different times in the summer of 2014. Considering the spectral range of APEX data (425-905 nm), macrophyte traits spatial variability maps are derived only for those spectral indices showing best matching with *in situ* measured data: i.e. NDSI_775,740_ as proxy of Chl-a content (R_cal_^2^ = 0.43), NDSI_433,665_ as proxy of Chl/Car ratio (R _cal_^2^ = 0.55), and NDSI_690,500_ as proxy of LMA (R _cal_^2^ = 0.44).

We fully recognise that the application of NDSI proxies using RS data inevitably bring to more or less significant biases, reflecting into the reliability of plant functional traits retrieved, which are due to complex combinations of factors, including: vegetation fractional cover and mixture with canopy background, density and structure effects (e.g. leaf orientation), reflectance anisotropy, and atmospheric effects (Asner *et al*., 2015). All the more so, spectral data measured from APEX airborne imager, with pixel size in this case of 5 m, are inherently measuring the response of macrophyte beds at canopy scale, and application of proxies derived and assessed at leaf scale to some extent hampers the absolute matching of observation scales for spectral reflectance and plant traits.

In spite of these distortion factors, APEX based maps of selected traits in Lake Hídvégi and Mantua lakes system, shown in Fig. 7, are clearly capturing notable patterns of relative variability in macrophyte communities at within-system scale that are informative about specific features of macrophytes growing the two sites. Indeed, spatial patterns of variability in spectral proxies for the three traits are highlighted in Fig. 7 maps and show differences at site scale, due to both inter- and intra-species differentiation (e.g. *Trapa* in Lake Inferior, *Nelumbo* in Lake Superior within Mantua lakes system, Fig. 7b), as well as considerable variability at community scale, indicating intra-specific trait plasticity (e.g. *Trapa* in Lake Hídvégi, Fig. 7a, and *Nelumbo* in Mantua lakes system, Fig. 7b). Differences in spatial patterns and ranges of trait proxies between and across the sites are not only due to species composition, but also to seasonal differences; temporal vegetation dynamics is actually another factor that can be taken under control by using remote spectroscopy data, once relations with plant traits are established. In fact, APEX data collected over Lake Hídvégi in middle of July represent macrophyte conditions at peak of growth, while data for Mantua lakes system, acquired in late September, provide a static image of macrophyte communities in early to late senescence phase, depending on species.

**Figure 7.**
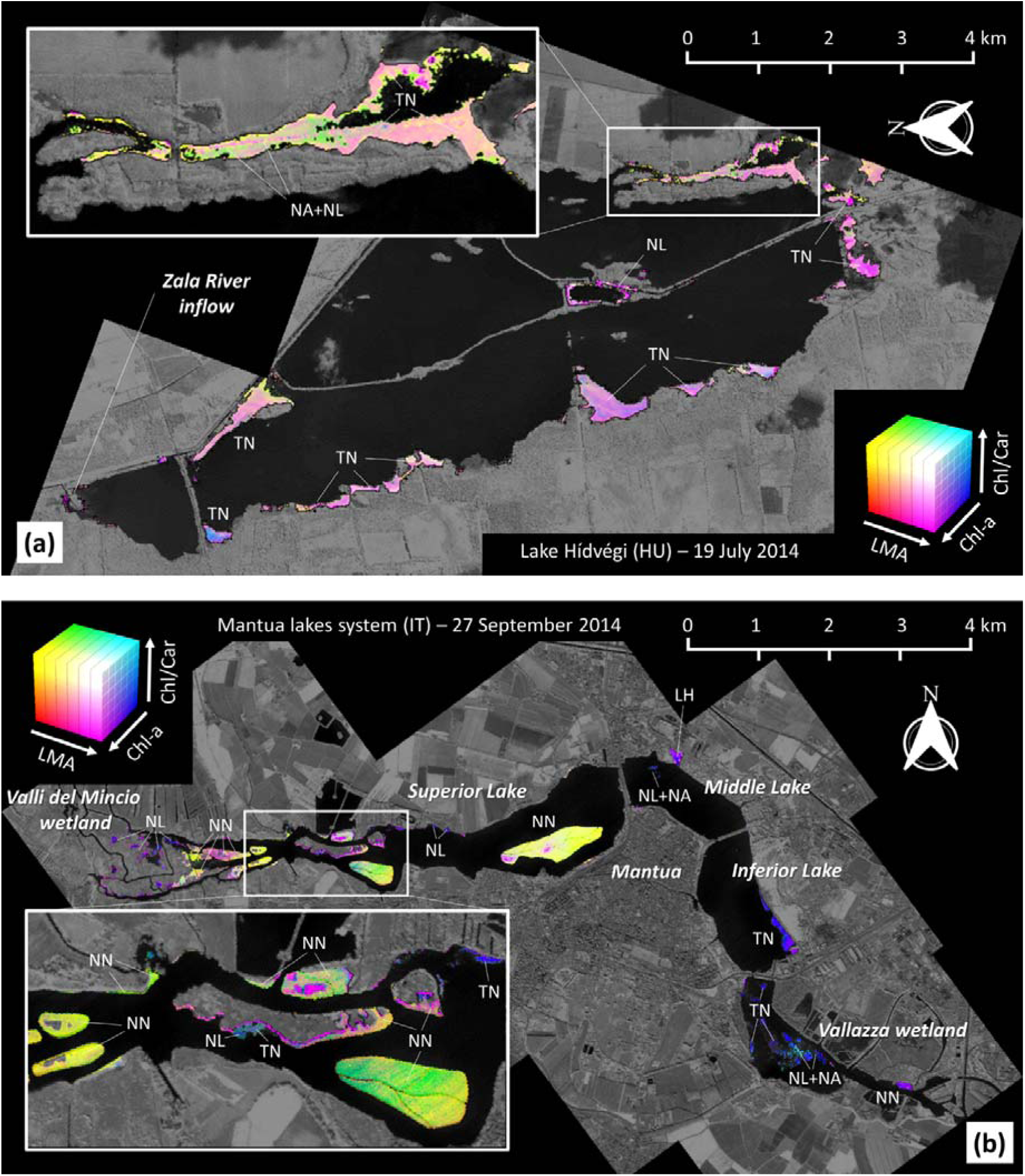
Maps of macrophyte trait spectral proxies derived from airborne hyperspectral (APEX) data over Lake Hídvégi (a) and Mantua lakes system (b), showing RGB combinations of best spectral proxies in the range covered by APEX data for: Chl-a content (NDSI_740,775_), Chl/Car ratio (NDSI_433,665_), and LMA (NDSI_690,500_). LH: *Ludwigia hexapetala*; NN: *Nelumbo nucifera*; NL: *Nuphar lutea*; NA: *Nymphaea alba*; TA: *Trapa natans*.

## 4. Discussion

For the specificity of their habitat, macrophytes can tolerate significant changes in environmental conditions during the growing season and across years, which may lead to alterations in physiological performance and community composition (Søndergaard *et al*., 2010). Macrophyte leaf trait data collected – covering different sites (three shallow lakes and wetlands in Europe), times (late May to late July), and seasons (three years) – showed a high degree of heterogeneity, both within and among species sampled. Allochthonous, invasive species tended to be photochemically different to native counterparts for most of the traits studied, while for some other traits all the species sampled were similar. In parallel with what observed for photophysiology, leaf structural traits of allochthonous species were found significantly different from those of autochthonous species. Differently from what found on mass basis in previous works (Tóth *et al*., 2019), foliar pigment content on area basis did not separate alien and native plants as distinctively as other traits, although a certain degree of segmentation at species level was found for pigments balance measures. This could imply that interspecific differentiation in studied macrophytes is driven more by pigment proportions, and therefore by the ability of species to adapt to given environment and maximise both light absorption and photoprotection (Grimshaw *et al*., 2002; Hussner *et al*., 2010). Such a multifaceted response, including both general and unique attributes of different species groups (i.e. allochthonous and autochthonous), not only shows the wide spectrum of macrophyte functional plasticity, but also hints at the individualistic responses of macrophytes to physical, chemical and anthropogenic characteristics of their environment.

Substantive intraspecific variability was found in our dataset, thus strengthening what already observed over Mantua lakes system in 2016-2017 by Tóth *et al*. (2019). These data indicate that high variability in both economical and photophysiological traits aids the studied macrophyte populations in expanding and persisting within their environment. Different species vary considerably in the lifespan of their leaves, resulting in increased foliar phenotypic plasticity and making direct functional interspecific comparison challenging, especially when effects of other environmental factors are to be considered (Kunii & Aramaki, 1992). Actual leaf age should therefore be taken into account for a thorough assessment of foliar trait variability among species, and it will be a key topic for future works.

Leaf reflectance is affected by multiple characteristics, with tri-dimensional leaf structure being the dominant factor, as outer (e.g. trichomes, cuticle surface) and inner (e.g. amount and ratio of different mesophylls) elements are responsible for major scattering mechanisms (Sims & Gamon, 2002; Klančnik & Gaberččik, 2016). Light absorption too is dependent on leaf structural arrangements, as more chlorophyll molecules are usually found in the palisade parenchyma, i.e. closer to the adaxial surface of the leaves (Terashima & Hikosaka, 1995; Smith *et al*., 1997), particularly when high porosity - the so-called aerenchyma in aquatic plants – is found (Borsuk & Brodersen, 2019). Our key finding is that a suite of leaf functional traits, connected with light absorption and scattering mechanisms and representative of foliar economics trade-off, can be effectively modelled across floating and emergent macrophyte species based on leaf reflectance, even under moderate to strong environmental heterogeneity, thus complementing works performed in terrestrial ecosystems (Homolová *et al*., 2013; Asner *et al*., 2015; Kattenborn *et al*., 2018; Van Cleemput *et al*., 2018).

Best results were obtained via PLSR models for structural traits (leaf DMC and LMA), which constitute synthetic descriptors of leaf morphology, and in particular the inner structure, mediating the effects of scattering mechanisms with water absorption. Maximum sensitivity of NDSI to DMC was found in SWIR range for two band combinations located around the shoulder of water absorption features (Kokaly *et al*., 2009), corroborated by VIP scores of PLSR models peaking around 1390-1400 nm (Fig. 6); overall modelling accuracy for DMC was higher than what recently reported for grassland species based on VNIR reflectance only (Wayne Polley *et al*., 2020). Leaf reflectance-LMA connections revealed a more complex picture, with optimal NDSI for macrophyte species sampled found for NDSI_1415,2305_ (R _cal_^2^ = 0.77), and low scores of VIP of PLSR model for wavelengths shorter than 1350 nm and between 1900 and 2100 nm. These patterns generally confirm what documented from previous works on terrestrial plants at macro spectral ranges, i.e. in the SWIR (le Maire *et al*., 2008; Féret *et al*., 2011; Wang & Li, 2012; Ali *et al*., 2017; Féret et al:, 2019), but with specific differences in optimal wavelengths to be used, possibly because of leaf structural peculiarities of aquatic plants (Klančnik & Gaberččik, 2016).

Complementing leaf structural traits in describing the trade-off at the core of the LES concept, leaf reflectance measured on the adaxial side of leaves proved to be a good predictor of biochemistry-related traits in macrophytes, represented here by pigments pools and ratios. Chl-a content and Chl/Car ratio were modelled with moderate to high reliability based on leaf spectroscopy and PLSR. Pigments are more directly related to spectral reflectance because of their primary function of interacting with light and their location within the leaf structure – i.e. in the first, adaxial strata aquatic plant leaves, as shown by Borsuk & Brodersen (2019) for chlorophylls in the case of *Eichornia crassipes* (Mart.) Solms. In general, relations found between leaf reflectance and pigments (Fig. 3), confirm and strengthen the notion that leaf spectral reflectance in the visible to NIR range, can be used as effective proxy for leaf pigments (and biochemistry) also in macrophytes. Optimal wavelengths for Chl-a predictions were found in the red-edge range (i.e. 700-800 nm), similar to what extensively documented for terrestrial plants (Gitelson *et al*., 2003; le Maire *et al*., 2008; Féret *et al*., 2011; Wang & Li, 2012), with VIP of PLSR model peaking around 705-710 nm. High NDSI-pigments correlations outside of PAR range, where pigments absorption is virtually null, can be attributed to interactions with other leaf traits (Supplementary Fig. S8), i.e. correlation with LMA for Car (*r* = +0.52) and Chl/Car (*r* = −0.48). Instead, best performing spectral combinations for Ca/Cb ratio found around 2200 nm is possibly driven by the link between Ca/Cb and N content (Kitajima & Hogan, 2003), which we did not measure and it is known to show absorption features around 2180 nm from previous works (Kokaly, 2001; Li *et al*., 2018).

The similarity between chlorophyll-a fluorescence dynamics and growth data suggests a close mechanistic link between photophysiological features and actual metabolism of plants (Genty *et al*., 1989; Peterson *et al*., 2001). Compared to what highlighted for leaf economics traits, reflectance-based PLSR models for photophysiological parameters were found to be under-performing across the species sampled. Best results were scored for F_v_/F_m_ (R_CV_^2^ = 0.21), with optimal spectral range very similar to photochemical reflectance index (PRI) linked to light use efficiency and xanthophyll cycle (Harris *et al*., 2014), and qP (R_CV_^2^ = 0.33), whose best spectral proxies were found in the SWIR range, roughly around 1400-1500 nm, probably due to co-variation of qP with DMC (*r* = +0.32). Sensitivity of chlorophyll fluorescence measures even to moderate environmental variations, paired with the large spatial-temporal variability typical of macrophyte populations - in the form of either trait variability within monospecific stands or specific trait distributions within mixed communities (Thomaz et al., 2009; Bornette & Puijalon, 2011) - and represented into our empirical dataset (Fig. 1; Table 1; Supplementary Table S1), could constitute a relevant obstacle to the establishment of general reflectance-photophysiology traits models, requiring specific spectral models limited to species and/or phenological stages (Supplementary Fig. S2-S7), as done by Stratoulias & Tóth (2020) for *Phragmites*.

Summarizing, our results demonstrate that spectral reflectance can be used for reliably estimating specific macrophyte leaf traits (Chl-a, Chl/Car, DMC and LMA), which are strongly connected to variability expressed within the LES in terms of trade-off between structural investment and photosynthetic efficiency (Poorter *et al*., 2009; Onoda *et al*., 2017), by using normalized spectral indices or PLSR modelling based on an empirical dataset collected across sites, seasons, and species. This finding provide the basis for using leaf spectra as a surrogate for high-throughput assessment of variability in macrophyte traits over scales and gradients, and support the extension of the reflectance-based trait modelling to RS spectroscopic data for enhancing the level of detail of functional diversity analysis. In particular, the possibility of using spectral proxies for modelling macrophyte LMA - a key trait in the LES as well as biodiversity variable (Pettorelli *et al*., 2016) - opens intriguing perspectives in aquatic species diversity and functioning, for future research and ecological applications of RS, aiming at large scale and investigating spatial and temporal gradients. The approach could be even extended to plant science in general, when paired with works already performed on terrestrial species (Serbin *et al*., 2019).

Spatial-wise maps of macrophyte traits as the ones derived from airborne hyperspectral data (Section 3.4) can provide intuitive, realistic and detailed *bio-visualisation* of vegetation diversity and connected processes. Indeed, the heterogeneity highlighted in Fig. 7 for macrophyte communities of Mantua lakes system and Lake Hídvégi could be hardly seen with punctual measurements of macrophyte traits, and yet it is captured in systematic, synoptic and quantitative way by applying leaf reflectance-traits relations we found to remotely sensed imaging spectroscopy data, informing us on spatial and temporal functional variability. Although some interesting case studies in this respect have been recently documented (e.g. Bolpagni *et al*., 2014; Tóth, 2018; Villa *et al*., 2018), they were limited to one or few systems and few aquatic vegetation characteristics (functional types, phenology metrics) and wider applications are needed.

Remote sensing spectroscopy can allow the quantitative assessment of changes in vegetation function across plant communities - at inter- and intra-specific levels - and this work provides the basis for developing research applied to inland water systems and connected ecotones, almost neglected in functional studies largely biased toward terrestrial ecosystems (Gustafsson & Norkko, 2017), tackling hot environmental issues such as dynamics of aquatic ecosystem under climate change and anthropogenic influence, as well as mechanisms of invasions by alien species.

## Acknowledgements

The authors thank the Parco del Mincio authority and voluntary ecological guards for support given during fieldwork in the Mantua lakes system. Many thanks to Mauro Musanti (CNR-IREA) and Attila Kovács (BLI, Centre for Ecological Research) for their help during fieldwork on Lake Hídvégi.

This work was supported by the agreement on scientific cooperation between the Hungarian Academy of Sciences and the Consiglio Nazionale delle Ricerche (MacroSense project), and by the Hungarian National Research, Development and Innovation Office, NKFIH [Grant No. KH-129505]. Part of the work was carried out in the context of the EU FP7 programme through the INFORM project [Grant No. 730066].

## Data accessibility

Leaf traits data will be deposited in Dryad (datadryad.org). Leaf reflectance spectra will be deposited also in EcoSIS (ecosis.org).

## Author contributions

PV, VRT designed the study, collected the data, performed the analysis, and drafted the manuscript. RB, MP supported collection of data and interpretation of results. All authors reviewed and edited the final manuscript.

