## Supplementary materials for "Leaf reflectance can surrogate foliar economics better than physiological traits across macrophyte species"

### *Supplementary Material*

**Supplementary Table S1.** Summary of *in situ* samples collected for this study.

| Location | | | | | Number of leaves sampled | | | |
|---|---|---|---|---|---|---|---|---|
| **Date** | **Site** | **Lat (N)** | **Lon (E)** | **Species** | **Reflectance spectra** | **Photophysiology traits** | **Pigments** | **Economics traits** |
| 21/07/2016 | Hídvégi | 46.6524 | 17.1433 | *Trapa* | - | - | 3 | 3 |
| 21/07/2016 | Hídvégi | 46.6581 | 17.1231 | *Trapa* | - | - | 3 | 3 |
| 21/07/2016 | Hídvégi | 46.6589 | 17.1238 | *Trapa* | - | - | 3 | 3 |
| 21/07/2016 | Hídvégi | 46.6131 | 17.1674 | *Trapa* | - | - | 3 | 3 |
| 22/07/2016 | Hídvégi | 46.6045 | 17.1665 | *Trapa* | - | - | 3 | 3 |
| 22/07/2016 | Hídvégi | 46.6003 | 17.1594 | *Trapa* | - | - | 3 | 3 |
| 22/07/2016 | Hídvégi | 46.6077 | 17.1428 | *Trapa* | - | - | 3 | 3 |
| 22/07/2016 | Hídvégi | 46.6135 | 17.1406 | *Trapa* | - | - | 2 | 2 |
| 21/07/2016 | Hídvégi | 46.6152 | 17.1675 | *Nuphar* | - | - | 3 | 3 |
| 21/07/2016 | Hídvégi | 46.6134 | 17.1676 | *Nymphaea* | - | - | 3 | 3 |
| 21/07/2016 | Hídvégi | 46.6146 | 17.1673 | *Nymphaea* | - | - | 3 | 3 |
| 27/07/2016 | Mantua | 45.1578 | 10.7463 | *Nelumbo* | 6 | 6 | - | - |
| 27/07/2016 | Mantua | 45.1608 | 10.7345 | *Nuphar* | 6 | 6 | - | - |
| 27/07/2016 | Mantua | 45.1601 | 10.7294 | *Phragmites* | 6 | 6 | - | - |
| 27/07/2016 | Mantua | 45.1608 | 10.7357 | *Trapa* | 5 | 5 | - | - |
| 27/07/2016 | Mantua | 45.1611 | 10.7331 | *Trapa* | 6 | 6 | - | - |
| 28/07/2016 | Mantua | 45.1624 | 10.7100 | *Ludwigia* | - | 6 | - | - |
| 28/07/2016 | Mantua | 45.1578 | 10.7141 | *Phragmites* | - | 6 | - | - |
| 29/07/2016 | Mantua | 45.1709 | 10.7971 | *Ludwigia* | - | 3 | - | - |
| 29/07/2016 | Mantua | 45.1643 | 10.8040 | *Trapa* | 6 | 6 | - | - |
| 29/07/2016 | Mantua | 45.1685 | 10.7929 | *Trapa* | 6 | 6 | - | - |

| | | | | | | | | |
|---|---|---|---|---|---|---|---|---|
| 29/07/2016 | Mantua | 45.1513 | 10.8121 | *Trapa* | 6 | 6 | - | - |
| 29/05/2017 | Mantua | 45.1624 | 10.7101 | *Ludwigia* | - | 12 | 9 | 9 |
| 29/05/2017 | Mantua | 45.1631 | 10.7819 | *Nelumbo* | 3 | 3 | - | - |
| 29/05/2017 | Mantua | 45.1629 | 10.7769 | *Nelumbo* | 3 | 3 | - | - |
| 29/05/2017 | Mantua | 45.1604 | 10.7670 | *Nelumbo* | 3 | 3 | - | - |
| 29/05/2017 | Mantua | 45.1625 | 10.7092 | *Nuphar* | - | 11 | 11 | 12 |
| 29/05/2017 | Mantua | 45.1608 | 10.7346 | *Nuphar* | 4 | 4 | 4 | 4 |
| 29/05/2017 | Mantua | 45.1689 | 10.7869 | *Nuphar* | 4 | 4 | - | - |
| 29/05/2017 | Mantua | 45.1606 | 10.7358 | *Trapa* | 4 | 4 | - | - |
| 30/05/2017 | Mantua | 45.1705 | 10.7929 | *Nymphaea* | 4 | 4 | 4 | 4 |
| 30/05/2017 | Mantua | 45.1647 | 10.8059 | *Trapa* | 4 | 4 | 4 | 4 |
| 30/05/2017 | Mantua | 45.1705 | 10.7929 | *Trapa* | 4 | 4 | 4 | 4 |
| 30/05/2017 | Mantua | 45.1686 | 10.7926 | *Trapa* | 4 | 4 | 4 | 4 |
| 30/05/2017 | Mantua | 45.1490 | 10.8146 | *Trapa* | 6 | 6 | 4 | 4 |
| 14/06/2017 | Hídvégi | 46.6040 | 17.1667 | *Trapa* | 6 | 6 | - | - |
| 14/06/2017 | Hídvégi | 46.6523 | 17.1427 | *Trapa* | 6 | 6 | - | - |
| 14/06/2017 | Hídvégi | 46.6584 | 17.1231 | *Trapa* | 6 | 6 | - | - |
| 14/06/2017 | Hídvégi | 46.6124 | 17.1414 | *Trapa* | 6 | 6 | - | - |
| 27/07/2017 | Mantua | 45.1627 | 10.7092 | *Ludwigia* | 7 | 9 | 7 | 9 |
| 27/07/2017 | Mantua | 45.1633 | 10.7828 | *Nelumbo* | 6 | 9 | 9 | 9 |
| 27/07/2017 | Mantua | 45.1623 | 10.7739 | *Nelumbo* | 6 | 6 | 6 | 6 |
| 27/07/2017 | Mantua | 45.1611 | 10.7682 | *Nelumbo* | 5 | 6 | 6 | 6 |
| 27/07/2017 | Mantua | 45.1627 | 10.7092 | *Nuphar* | 3 | 3 | 2 | 3 |
| 27/07/2017 | Mantua | 45.1633 | 10.7473 | *Trapa* | 9 | 9 | 7 | 8 |
| 28/07/2017 | Mantua | 45.1704 | 10.7921 | *Nuphar* | 6 | 5 | 6 | 6 |
| 28/07/2017 | Mantua | 45.1705 | 10.7927 | *Nymphaea* | 6 | 6 | 6 | 6 |
| 28/07/2017 | Mantua | 45.1688 | 10.7913 | *Trapa* | 6 | 6 | 6 | 6 |
| 28/07/2017 | Mantua | 45.1652 | 10.8053 | *Trapa* | 6 | 6 | 6 | |
| 28/07/2017 | Mantua | 45.1495 | 10.8142 | *Trapa* | 6 | 6 | 5 | 6 |
| 17/07/2018 | Hídvégi | 46.6149 | 17.1675 | *Nuphar* | 6 | 6 | - | - |
| 17/07/2018 | Hídvégi | 46.6146 | 17.1674 | *Nymphaea* | 6 | 6 | - | - |
| 17/07/2018 | Hídvégi | 46.6524 | 17.1431 | *Trapa* | 5 | 6 | - | - |
| 17/07/2018 | Hídvégi | 46.6588 | 17.1241 | *Trapa* | 5 | 6 | - | - |
| 18/07/2018 | Hídvégi | 46.6003 | 17.1601 | *Trapa* | 6 | 6 | - | - |
| 18/07/2018 | Hídvégi | 46.6127 | 17.1413 | *Trapa* | 6 | 6 | - | - |
| 24/07/2018 | Varese | 45.8065 | 8.7661 | *Ludwigia* | 12 | 12 | - | - |
| 24/07/2018 | Varese | 45.8068 | 8.7717 | *Ludwigia* | 12 | 12 | - | - |
| 24/07/2018 | Varese | 45.8144 | 8.7581 | *Nelumbo* | 6 | 6 | - | - |
| 24/07/2018 | Varese | 45.8138 | 8.7588 | *Nelumbo* | 6 | 6 | - | - |
| 24/07/2018 | Varese | 45.8042 | 8.7756 | *Trapa* | 6 | 6 | - | - |
| 25/07/2018 | Mantua | 45.1626 | 10.7092 | *Ludwigia* | 10 | 12 | 12 | 12 |
| 25/07/2018 | Mantua | 45.1626 | 10.7098 | *Ludwigia* | 12 | 12 | 11 | 12 |
| 25/07/2018 | Mantua | 45.1633 | 10.7838 | *Nelumbo* | 6 | 6 | 6 | 6 |
| 25/07/2018 | Mantua | 45.1614 | 10.7689 | *Nelumbo* | 5 | 6 | 6 | 6 |
| 25/07/2018 | Mantua | 45.1632 | 10.7475 | *Trapa* | 6 | 6 | 5 | 6 |
| 26/07/2018 | Mantua | 45.1704 | 10.7933 | *Nuphar* | 6 | 6 | 6 | 6 |
| 26/07/2018 | Mantua | 45.1704 | 10.7933 | *Nymphaea* | 6 | 6 | 6 | 6 |
| 26/07/2018 | Mantua | 45.1652 | 10.8050 | *Trapa* | 6 | 6 | 6 | 6 |
| 26/07/2018 | Mantua | 45.1686 | 10.7920 | *Trapa* | 6 | 6 | 6 | 6 |
| 26/07/2018 | Mantua | 45.1482 | 10.8148 | *Trapa* | 6 | 6 | 6 | 6 |

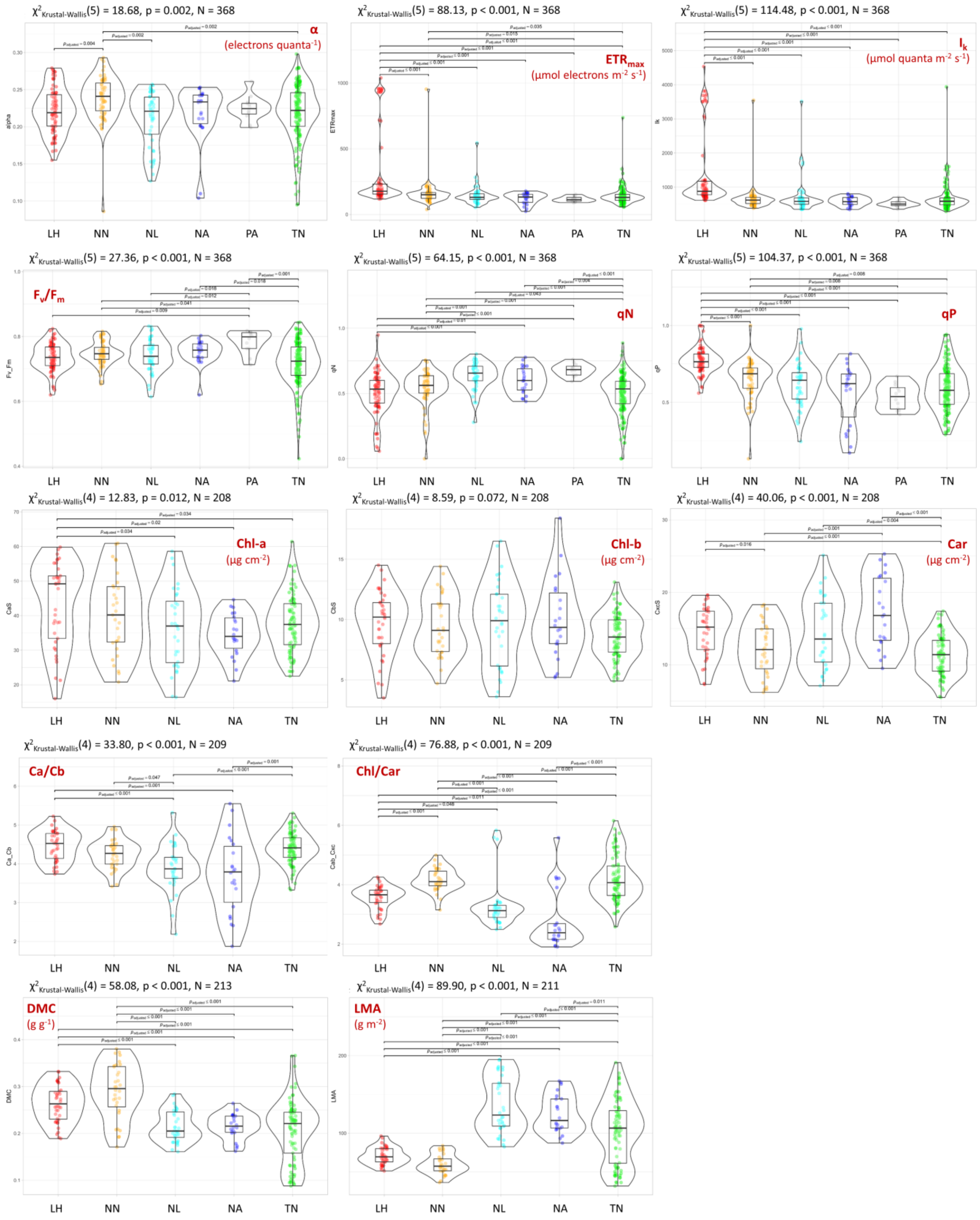


**Supplementary Figure S1.** Violin plots (with encompassed box plots) showing range and distribution of all leaf traits - photophysiological parameters, pigments content and leaf economics spectrum traits - measured over 6 macrophyte species (LH=*Ludwigia hexapetala*; NN=*Nelumbo nucifera*; NL=*Nuphar lutea*; NA=*Nymphaea alba*; PA=*Phragmites australis*; TN=*Trapa natans*). Plots show significant differences ($p < 0.05$) in pairwise comparison performed via Dunn's post-hoc test with Benjamini-Hochberg adjustment.

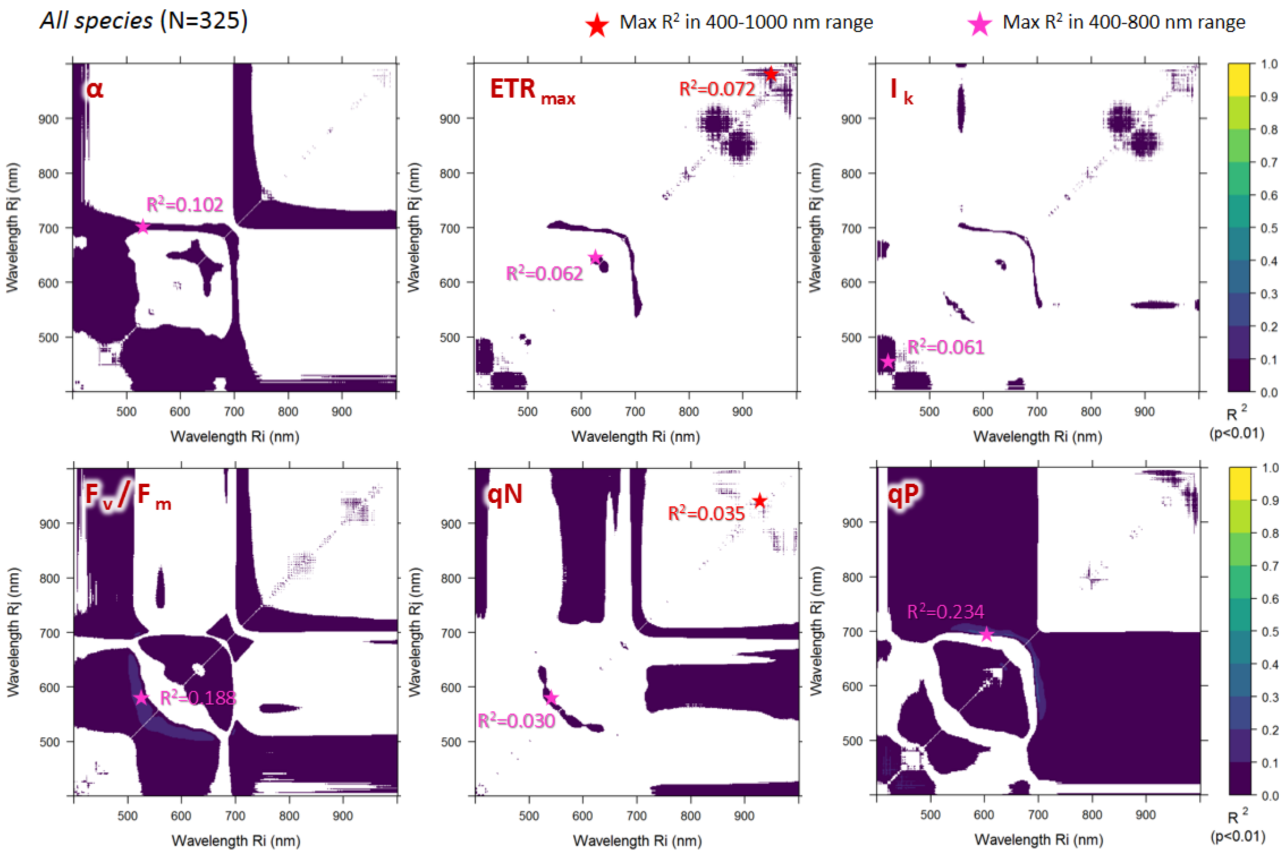


**Supplementary Figure S2**. Statistically significant (p < 0.01) NDSI correlations with photophysiological parameters measured on all macrophyte species sampled (N=324) in the visible to near-infrared spectral range (400-1000 nm).

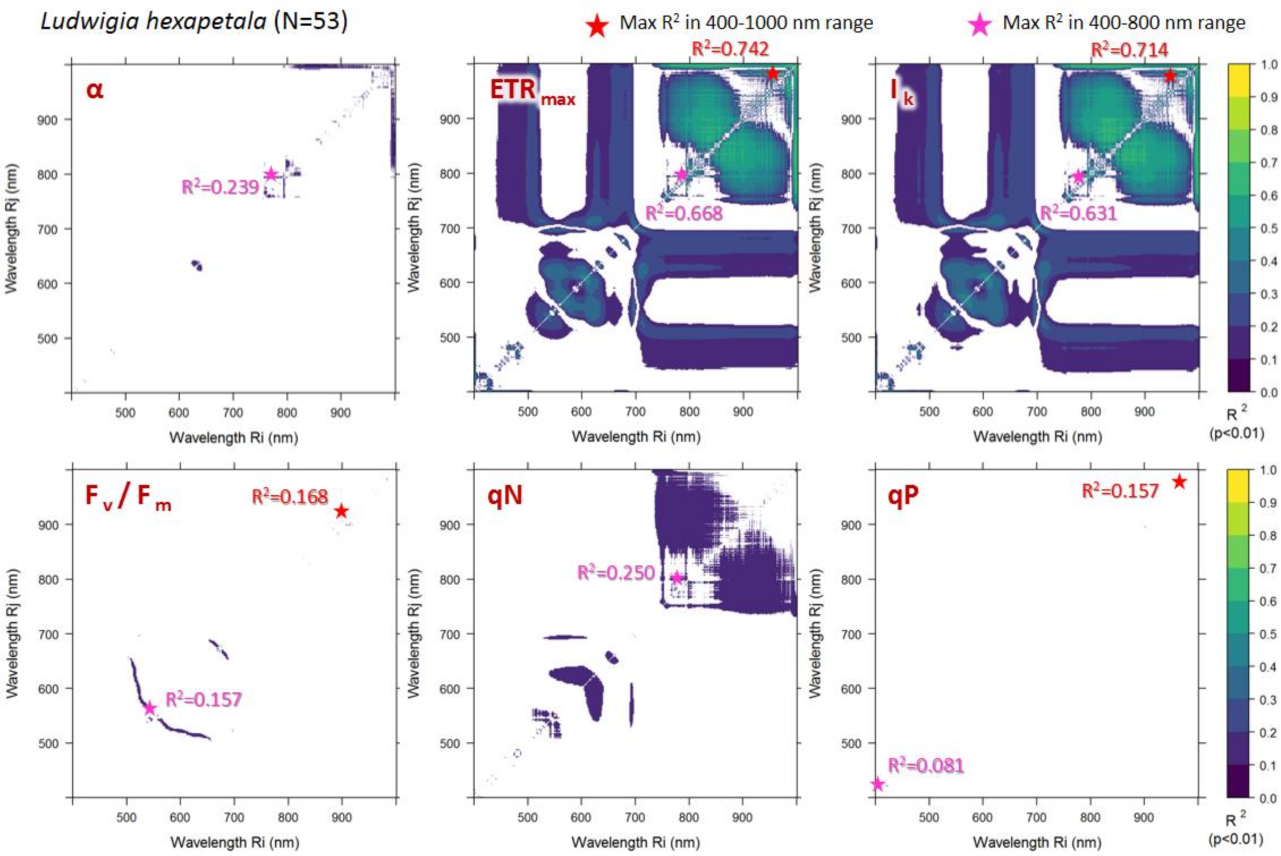


**Supplementary Figure S3**. Statistically significant (p < 0.01) NDSI correlations with photophysiological parameters measured on *Ludwigia hexapetala* samples (N=53) in the visible to near-infrared spectral range (400-1000 nm).

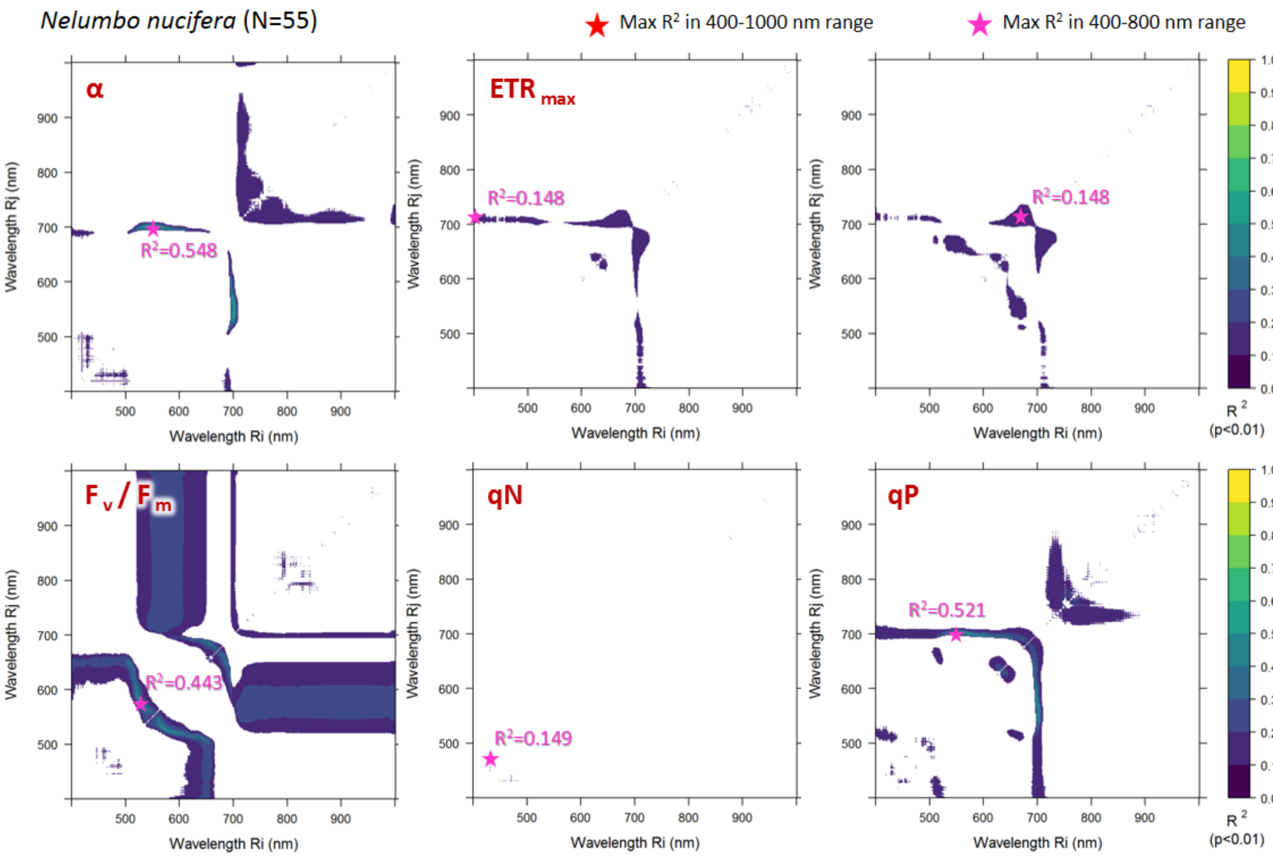


**Supplementary Figure S4**. Statistically significant ($p < 0.01$) NDSI correlations with photophysiological parameters measured on *Nelumbo nucifera* samples (N=55) in the visible to near-infrared spectral range (400-1000 nm).

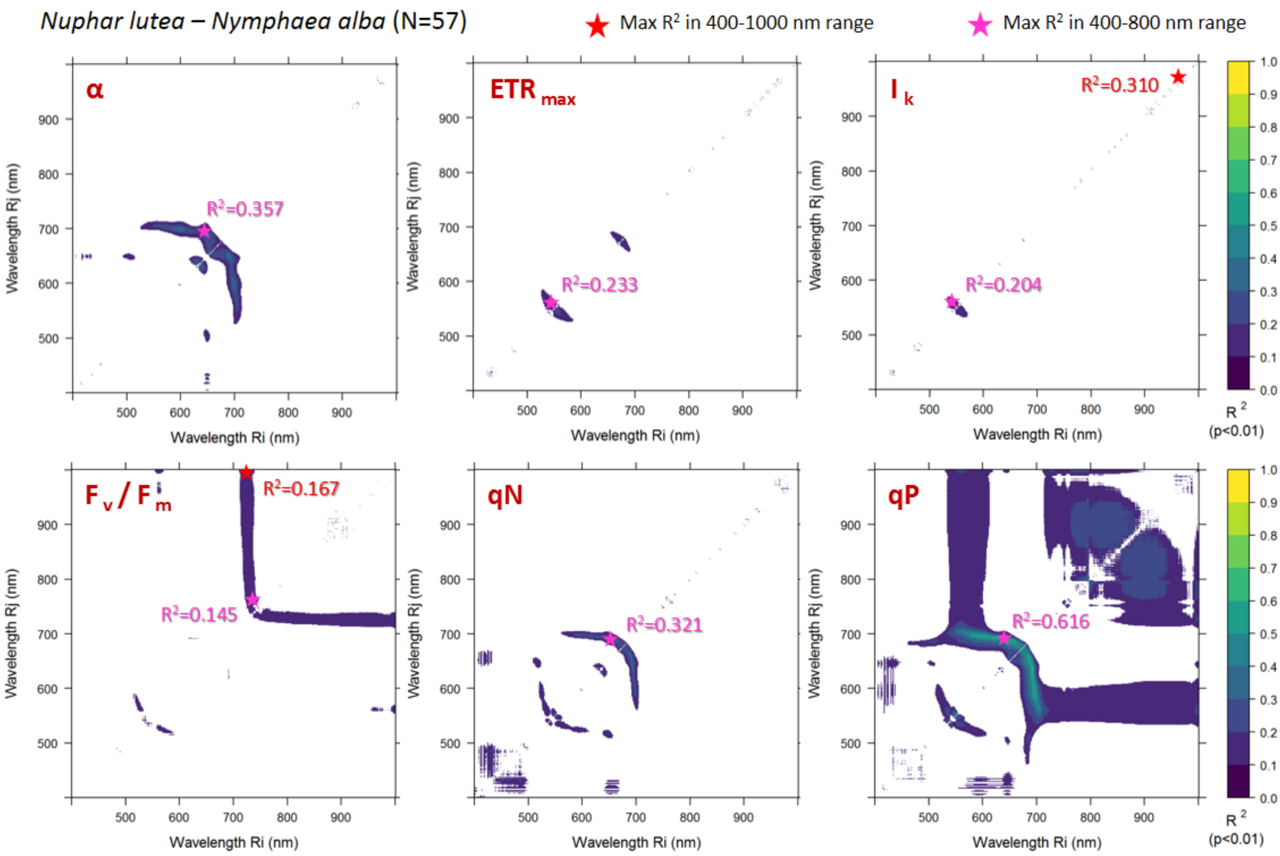


**Supplementary Figure S5**. Statistically significant ($p < 0.01$) NDSI correlations with photophysiological parameters measured on *Nuphar lutea* and *Nymphaea alba* samples (N=57) in the visible to near-infrared range (400-1000 nm).

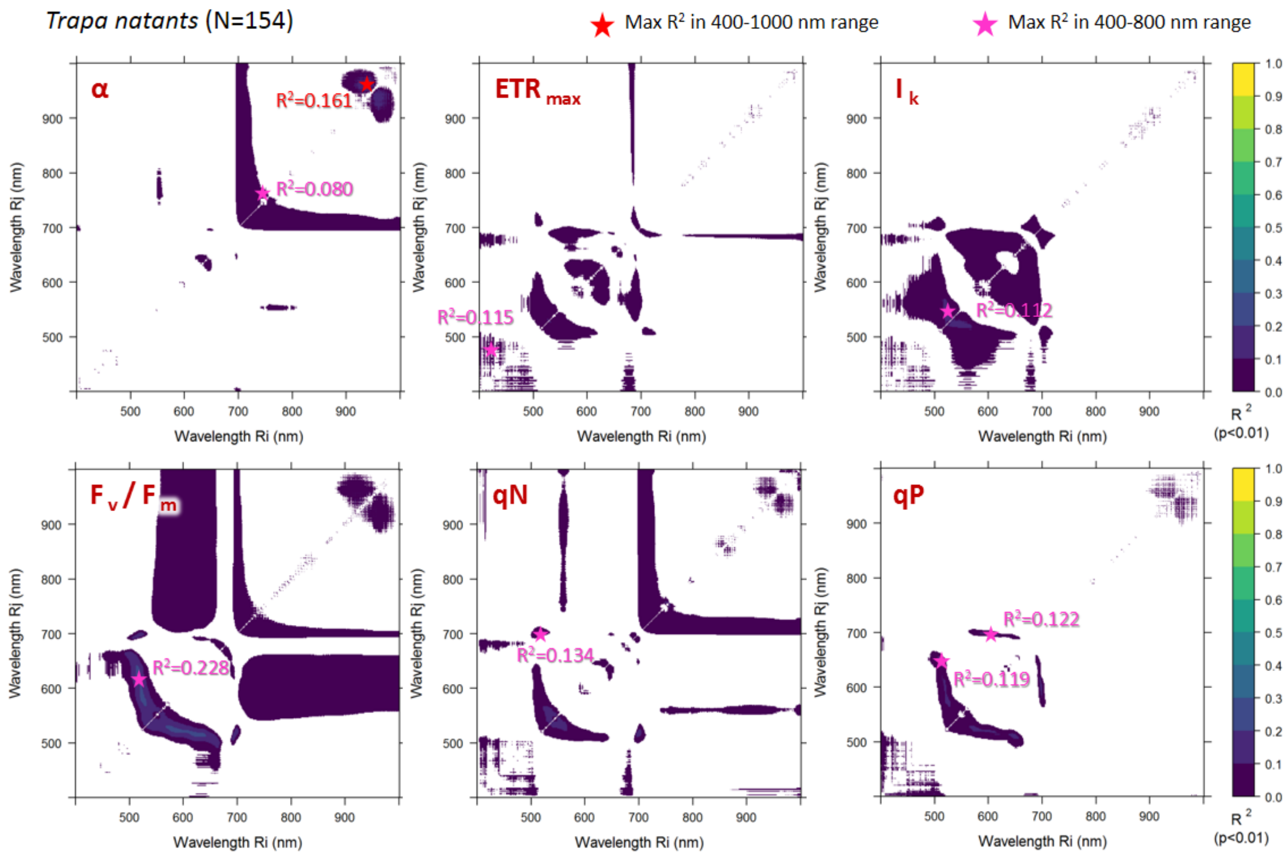


**Supplementary Figure S6**. Statistically significant ($p < 0.01$) NDSI correlations with photophysiological parameters measured on *Trapa natans* samples (N=154) in the visible to near-infrared spectral range (400-1000 nm).

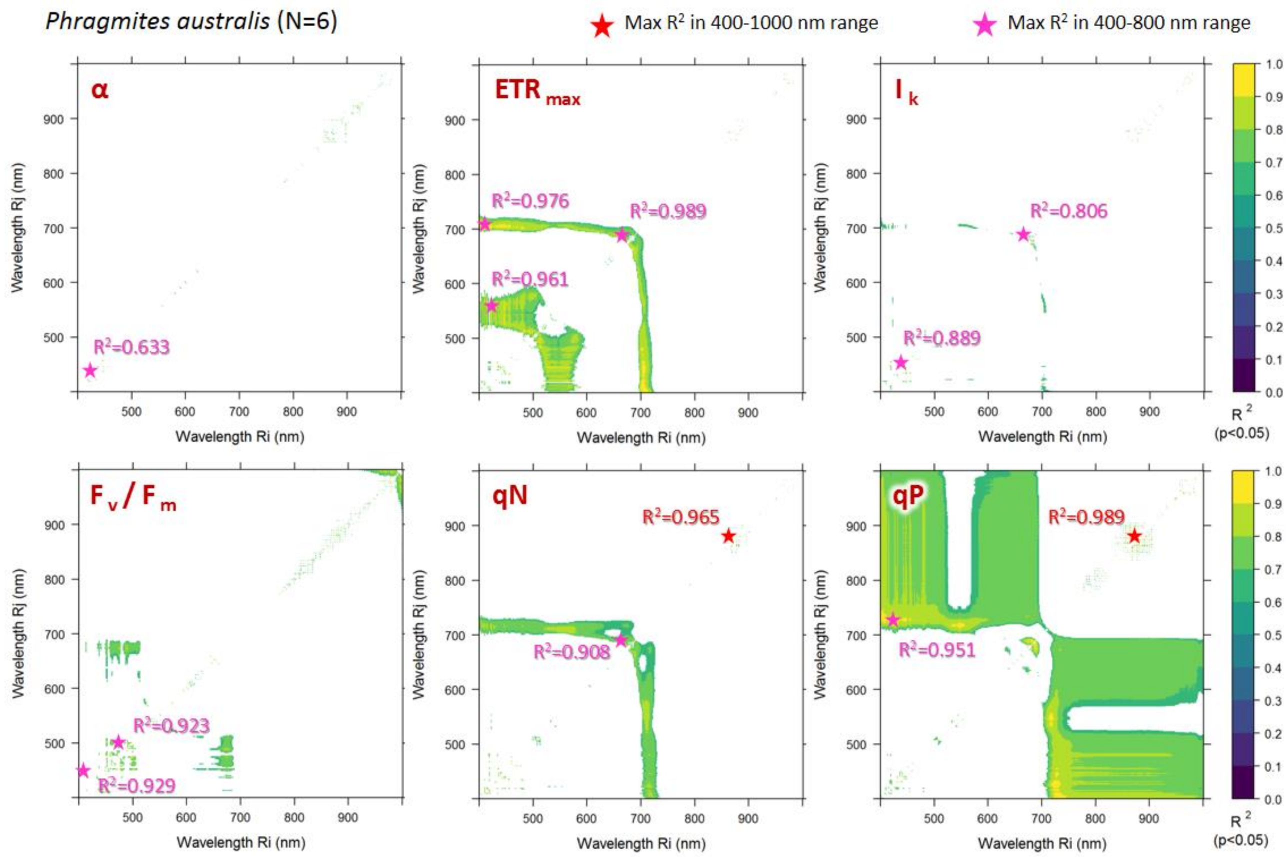


**Supplementary Figure S7**. Statistically significant ($p < 0.01$) NDSI correlations with photophysiological parameters measured on *Phragmites australis* samples (N=6) in the visible to near-infrared spectral range (400-1000 nm).

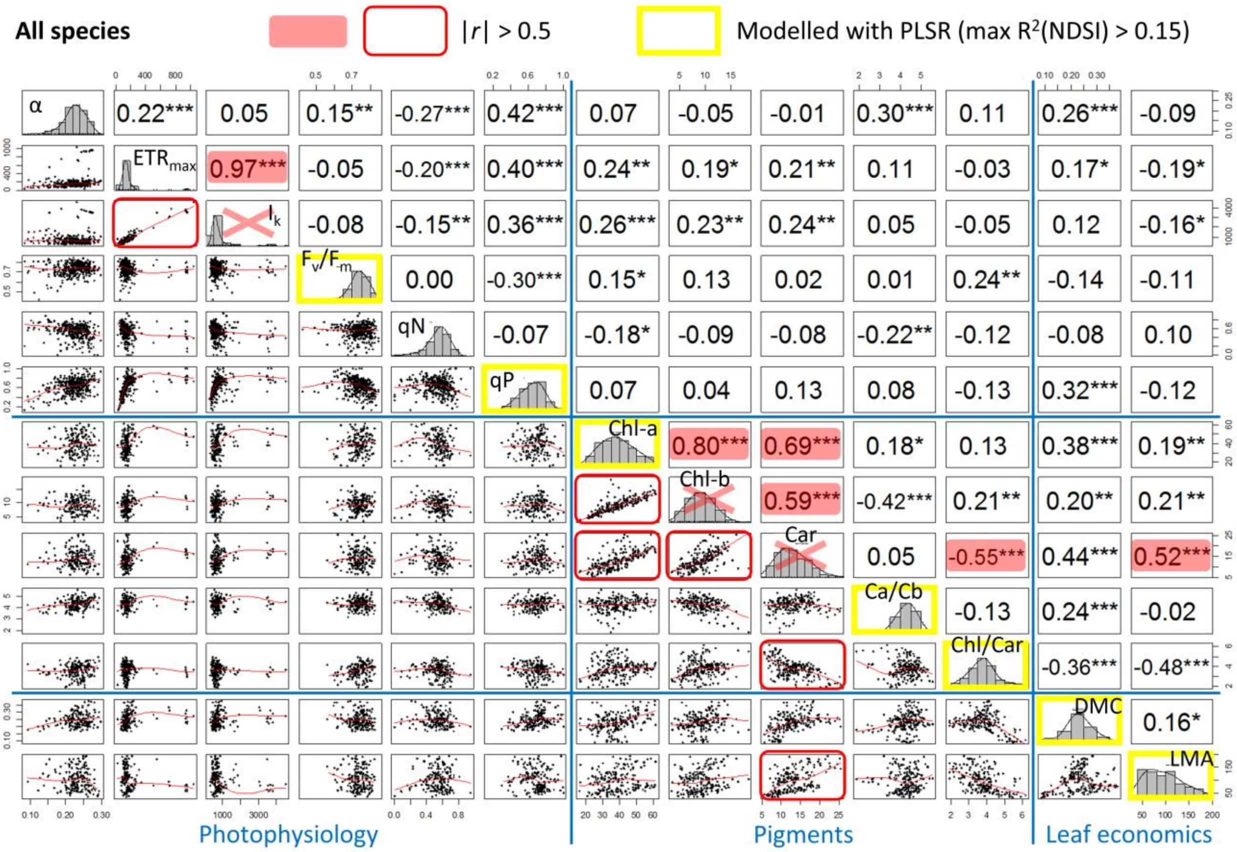


**Supplementary Figure S8.** Correlation matrix of leaf traits measured over 6 species. Pairwise scatter plots are shown in the lower left half, histograms are shown on the diagonal, and the coefficient of correlation (Pearson's r) of each pair of traits is shown in the upper right half, including info about its significance level (* p<0.05, ** p<0.01, *** p<0.001).

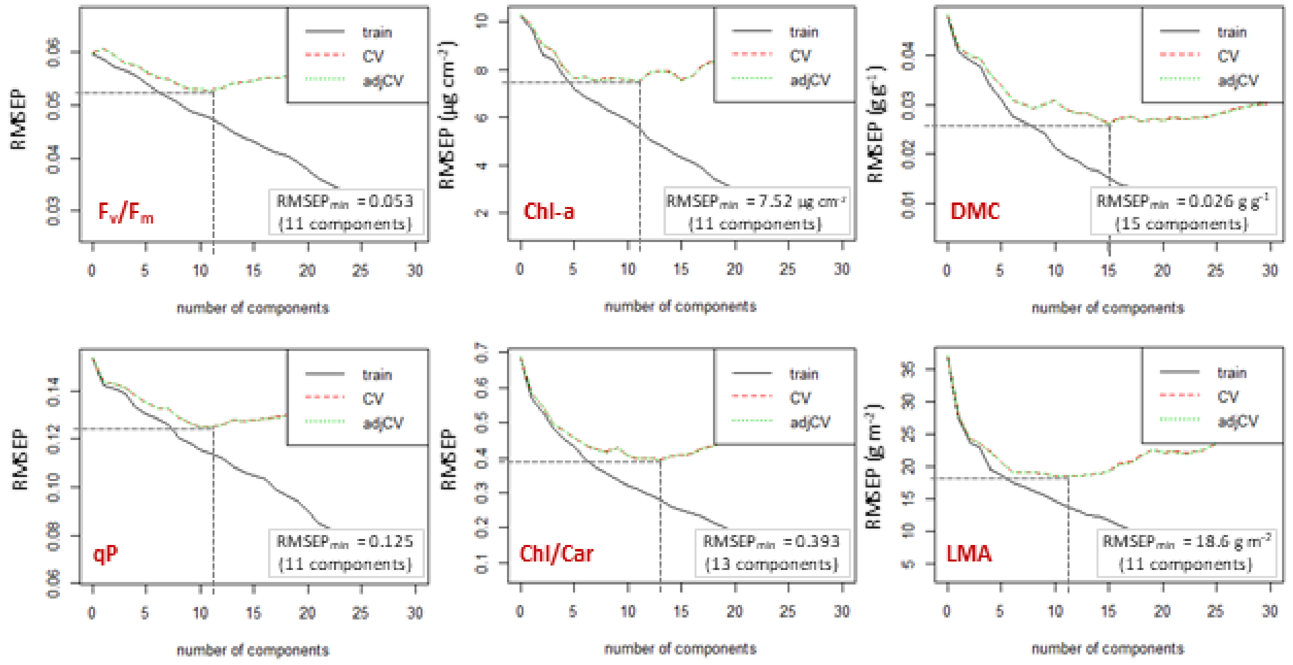


**Supplementary Figure S9.** Variation of root mean square error of prediction (RMSEP) with PLSR model components, computed against the full dataset (training as well as via leave-one-out cross-validation (CV) for key leaf traits modelled based on leaf spectral reflectance of measured macrophyte species.
